# A synthetic elastic protein as molecular prosthetic candidate to strengthen vascular wall elasticity

**DOI:** 10.1101/2023.11.30.569188

**Authors:** Marie Hoareau, Chloé Lorion, Lauriane Lecoq, Aurore Berthier, Baptiste Pierrat, Stéphane Avril, Fabrice Pirot, Pascal Sommer, Jérôme Sohier, Elise Lambert, Romain Debret

## Abstract

The loss of elasticity is a hallmark of systemic aging or genetic syndromes (e.g. cutis laxa, Williams-Beuren and supravalvular aortic stenosis) with direct consequences on tissue functions, and particularly deleterious when associated to the cardiovascular system. Tissue elasticity is mainly provided by large elastic fibers composed of supramolecular complexes of elastin and microfibrils. In arteries, the mature elastic fibers are located in the media compartment and form concentric elastic lamellar units together with the smooth muscle cells (SMCs). The main function of vascular elastic fibers is to allow extension and recoil of the vessel walls in response to the intraluminal pressure generated by the blood flow following cardiac systole. The synthesis of elastic fibers (elastogenesis) mainly occurs during the last third of fetal life with a peak in the perinatal period and then slowly decreases until the end of growth; as a result, elastic fiber repair is almost non-existent in adults. To date, no treatment exists to restore or repair deficient or degraded elastic fibers. A few pharmacological compounds have been proposed, but their efficacy/side effects balance remains very unfavorable. As an alternative strategy, we developed a synthetic elastic protein (SEP) inspired by the human tropoelastin, the elastin soluble precursor, to provide an elastic molecular prosthesis capable of integrating and reinforcing endogenous elastic fibers.

The SEP was easily produced in E. coli and purified by inversed transition cycling method. The resulting 55 kDa protein recapitulates the main physicochemical properties of the tropoelastin as thermal responsiveness, intrinsically disordered structures, and spherical self-assembly. The cross-linked SEP displays linear elastic mechanical properties under uniaxial tension loads. Using a co-culture *in vitro* model of the endothelial barrier, our results show that SEP is able to cross the cohesive endothelial monolayer to reach underlying SMCs. Moreover, SEP is processed by SMCs through a lysyl oxidase-dependent mechanism to form fibrillar structures that colocalize with fibrillin-rich microfibrils. The SEP was further characterized *in vivo* through the zebrafish model. The results indicate a global innocuity on zebrafish embryos and an absence of neutrophil recruitment following injection into the yolk sac of zebrafish. Finally, intravenous injection of a fluorescent SEP highlights its deposition in the wall of tortuous vessels which persists for several days after injection of the larvae. Taken together, our results demonstrate for the first time the incorporation of a naked tropoelastin-bioinspired polypeptide in endogenous elastic fibrillar deposits from SMCs, and its recognition by the lysyl-oxidase enzymatic machinery. In absence of toxicity and proinflammatory signal combined to a long-lasting accumulation in vessels *in vivo*, the SEP fulfills the first prerequisites for the development of an original biotherapeutic compound addressing the repair of elastic fibers.

## Introduction

Elasticity is essential to ensure the proper functioning of multiple organs and tissues such as skin, lungs, blood vessels, tendons, and elastic cartilages. Elastic fibers are the main source of elasticity in the body. These fibers are composed of two major proteins: tropoelastin and fibrillin. Tropoelastin monomers first assemble at the surface of the cell to form small aggregates, which are then reticulated on the fibrillin-rich microfibril scaffold to form the hydrophobic elastin core and finally the elastic fiber [1–3]. Other proteins are necessary for the correct assembly of the fibers such as fibulins and microfibril-associated glycoproteins [4], and their maturation by specific cross-linking network orchestrated by lysyl oxidases (LOXs) [5].

Elastic fibers are mostly synthesized perinatally and their formation is strongly downregulated after birth [6]. Healthy tissues do not undergo significant renewal of their fibers, meaning that they have to stay stable and functional throughout life [7, 8]. Although elastin is quite resistant to enzymatic degradation [9], elastic fibers tend to accumulate damage with time. Elastin and elastic fibers are indeed subject to several age-related modifications such as mechanical fatigue, calcification, and chemical alterations (glycation, carbamylation and peroxidation). They are also directly susceptible to proteolysis by specific proteases and matrix-metalloproteinases (MMPs) [10]. All these age-related alterations weaken the fibers and can have important consequences on health [11]. The fragmentation of elastin also leads to the formation of elastin-derived peptides (EDPs), some of which are bioactive (sometimes called elastokines or matrikines) and able to modify cell behavior [12] with potential impacts on cell adhesion, proliferation, chemotaxis, protease activation and apoptosis for example [10].

Elasticity deficiencies can also result from acquired or inherited disorders (reviewed in [13]). Among them, some are directly linked to a defect in the tropoelastin gene, such as Williams Beuren syndrome (WBS) [14] and the autosomal dominant cutis laxa (ADCL) [15]. The most impacted tissues are skin, blood vessels and lungs with a broad range of symptoms. WBS is often associated with supravalvular aortic stenosis (SVAS), a narrowing of the aortic root [16]. Other arteries are also subject to stenosis in this pathology [17]. In ADCL, patients can suffer from aortic aneurism, pulmonary emphysema, hernia and high skin laxity [15]. Some cases of acquired cutis laxa have also been described, in which the elastic fibers are more susceptible to degradation during inflammatory responses [18].

It is clear that cardiovascular complications are the most deleterious symptoms associated with the loss of elastic fiber integrity. Elastic arteries, such as the aorta, carotid and pulmonary arteries, are particularly susceptible to the deterioration of the mechanical function because these structures, located in direct proximity to the heart, must absorb the significant intraluminal distending pressure following the ejection of blood during the systolic phase. Thus, a higher number of contractile-elastic lamellar units (comprising a layer of SMCs and a lamella of elastic fibers) is requested in proximal arteries and this number decreases as distally as the caliber of arteries diminishes. Moreover, the inner elastic lamellar units undergo wider extension compared to outer units to homogenize layer extension through the arterial wall [19]. Consequently, the number of broken lamellar units noted histologically is a hallmark of artery pathophysiology and disease progression [20]. Currently, there is no satisfactory treatment to prevent, cure or restore elastic fiber integrity. Minoxidil, a K^+^-channel opener, was shown to promote elastic fiber neosynthesis in aged mice [21] and to confer long-term improvement in end-organ perfusion potentially by remodeling the arterial matrix in *Eln*^+/-^ mice [22]. However, it is also associated with important side effects, including cardiac hypertrophy and edemas formation [21]. For severe cases of SVAS, a surgical intervention is often necessary, but it does not treat the underlying causes and the risk of reoperation or mortality remains high after surgery [23].

An interesting alternative to treat elastin deficiency would be to replace it directly in affected tissues with a molecule able to integrate elastic fibers and act as the human tropoelastin (hTE) *in vivo*. Elastin-like recombinamers (ELRs) are synthetic polypeptides bioinspired by the hTE. They are widely used in biomaterial engineering for their interesting physical and mechanical behavior and have been increasingly used in the biomedical field in the last few years [24]. They can be considered as smart polymers as they are able to respond to physicochemical parameters of their microenvironment and undergo changes in response to it. They conserve the interesting properties of the hTE, such as its high resilience upon stretching, its self-assembly capacity and its biocompatibility [25]. In addition, ELRs allow to precisely control their molecular weight, functionality, and mechanical properties by modifying their amino acid sequence at will. ELRs are now investigated for diverse biomedical applications, particularly in drug delivery [26], nanovaccines [27], but also as biocompatible stent-coating to improve the success of angioplasty [28, 29], or more recently for the engineering of venous valves [30] just to cite a few.

As a matter of fact, products inspired by elastin are exclusively used as building blocks in the formulation of biomaterials, either a vehicle or a scaffold for tissue engineering. In the present study, we have designed a new ELR for biotherapeutic purposes with the ambition to deliver an agent that can be directly processed by cells to strengthen elastic fibers. We therefore produced and purified a recombinant synthetic elastic protein (SEP) that we characterized using physicochemical and mechanical analysis tools. Then we observed how this SEP was tolerated and used by human vascular cells (endothelial and smooth muscle cells). Finally, we administered the SEP in zebrafish models to determine its biocompatibility and its distribution in the vessels over time.

## Results

### Production and formulation

An original synthetic elastic protein (SEP) (Fig. 1A) was designed to conform with several remarkable specificities of the human tropoelastin (hTE). This includes alternative hydrophobic (VGVAPG-VGVLPG) and crosslinking (AAAKAAAKAAK) domains, together with the presence of the C-terminal domain 36 of the hTE since it was recognized to facilitate elastic fibers assembly through the GRKRK motif and the disulfide bond [31]. The choice of primary sequences has been based on short repeated motifs that are conserved across species to avoid exacerbate immunogenic responses. In addition, a Flag tag octapeptide was added in the N-terminal region to facilitate SEP recognition and discrimination from the native hTE using specific anti-Flag antibodies. A synthetic gene coding for the SEP was inserted in the pET30a procaryotic expression vector to transform the BL21(DE3) bacterial strain. After fermentation process and cell lysis, SEP was found in equivalent amount in both soluble and insoluble phases indicating that part of the production is sequestered in inclusion bodies of the bacteria probably due to its intrinsic hydrophobicity. Its electrophoretic profile showed apparent molecular weights of 55 kDa and 110 kDa for the monomer and a probable dimer form of the protein respectively (Fig. 1B). The SEP contained in the soluble phase was then subjected to inverse transition cycling (ITC) purification based on the method described by Meyer and Chilkoti [32]. One to three cycles were usually effective to discard bacterial material and obtain the SEP with high purity grade (Fig. 1C).

**Fig. 1.**
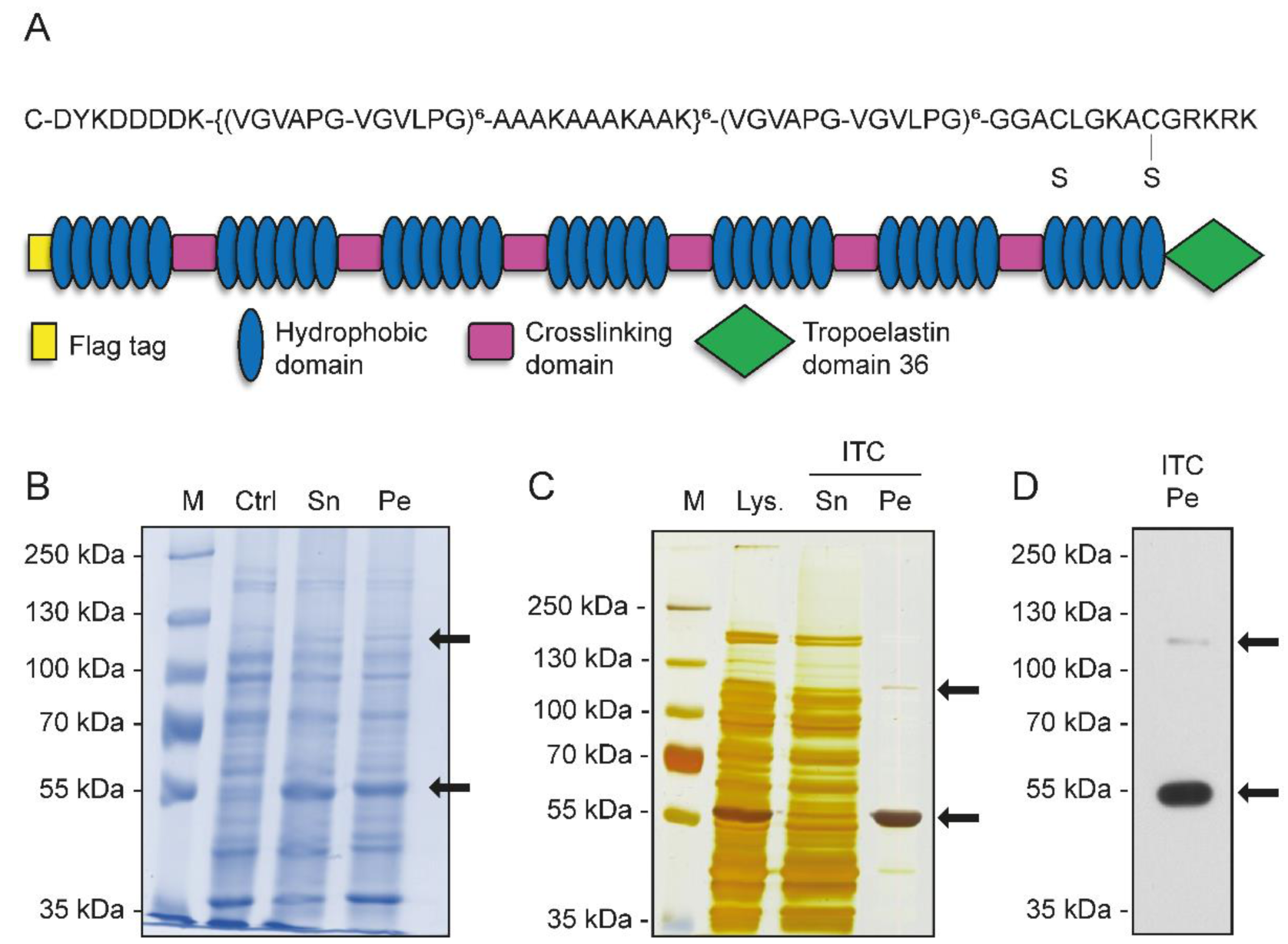
SEP is produced in E. coli and is easily purified by ITC. **A)** Schematic representation of SEP and its primary sequence. **B)** SEP over-expression. Cell lysate was analyzed by SDS-PAGE stained with Coomassie blue. Results are shown for pre-induced total cell lysate (Ctrl) and induced culture phases (Sn = supernatant, Pe = pellet). Black arrows indicate SEP monomer and dimer. **C)** SEP purification by inverse transition cycling (ITC). Total cell lysate (Lys), soluble fraction (Sn) and the pellet (Pe) containing SEP were analyzed by SDS-PAGE and the gel was stained by silver nitrate. **D)** SEP identification by western blot analysis using an anti-FLAG antibody.

Identification of the SEP was performed by western blot using an anti-Flag antibody and the two forms (monomeric and dimeric) were confirmed (Fig. 1D).

SEP was formulated in aqueous solution at 1 or 10 mg/mL in 0.9 % (w/v) NaCl. Endotoxin units were measured at 5.179 IU/mL for the 1 mg/mL formulation, which is below the 350 IU/mL limit required for biomedical applications. Solution osmoses were respectively measured at 288 mOsm and 315 mOsm for the 1 mg/mL and 10 mg/mL preparations, allowing these formulations to be injected in the blood flow with very low hemolytic risk.

### Structure characterization

Circular dichroism spectroscopy was used to determine secondary structures of SEP. Spectra were recorded at 0.2 mg/mL in phosphate buffer with increasing temperature (Fig. 2A). From 5°C to 25°C, the profile of SEP was characterized by a deep negative peak at 198 nm with a shoulder at 222 nm, which is indicative of a poorly ordered structure, mostly made of random coiled motifs, with the likely presence of a minority α-helix. When the temperature was raised above 35°C, the 222 nm peak was slightly red-shifted, resulting from a probable decrease of helix stability.

**Fig. 2.**
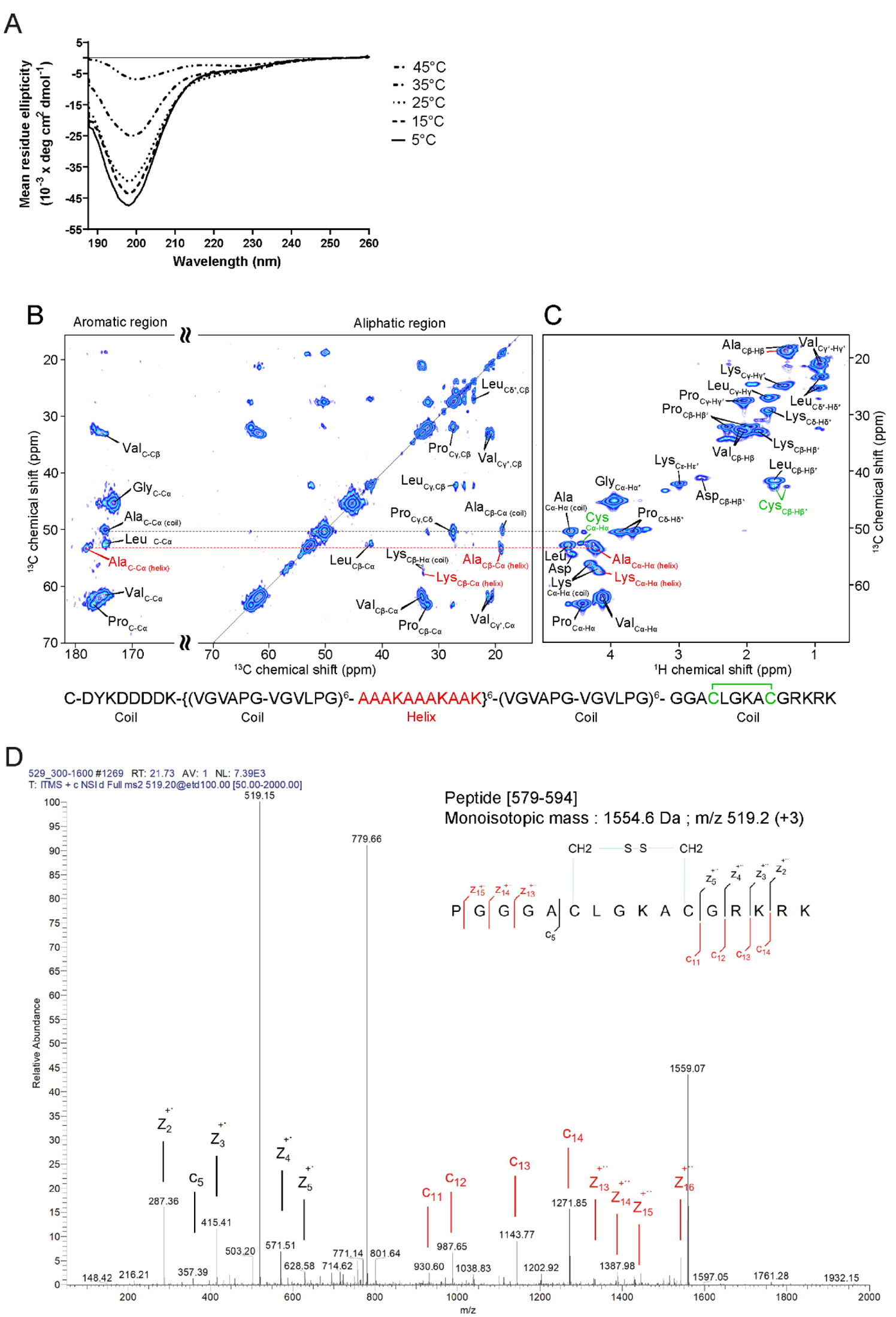
SEP displays secondary structures and a disulfide bound. **A)** Temperature-dependent CD spectra recorded in PBS pH 7,4 at 5°C, 15°C, 25°C, 35°C and 45°C. **B)** DARR and **C)** INEPT NMR spectra of SEP coacervates. **D)** LC-MS/MS analysis: fragmentation of the SEP peptide [579-

NMR spectra were recorded to gain conformation information on a residue-type basis (Fig. 2B- C). The 2D CC DARR spectrum allows to detect the rigid parts of the protein (Fig. 2B), while the 2D HC INEPT spectrum reveals the flexible parts (Fig. 2C). Globally, SEP is a flexible molecule since all residue types are detected in the INEPT spectrum. Residue types can be assigned based on their chemical shifts, revealing that all residues could be identified in the spectra, except the N-terminus Tyr which is present in lower amount compared to other residues for which repetitive motifs are present. All residue types identified are in coil conformation according to their chemical shifts (Asp, Val, Gly, Ala, Pro and Leu). In addition, two residue types are identified in the helical conformation, namely Ala and Lys as seen in Fig. 2B. These results are in accordance with CD data, and allow to confirm the α-helix structure in the region (AAAKAAAKAAK). Furthermore, cysteines are detected in the INEPT spectrum at a chemical shift (Cα = 42.7 ppm and Cβ = 52.6 ppm) typical of the oxidized form [33], revealing the presence of a disulfide bound between the two cysteines in the C-ter region of SEP (Fig. S1).

Finally, SEP was subjected to liquid chromatography tandem mass spectrometry (LC-MS/MS) in electron transfer dissociation (ETD) mode. The resulting spectra focused on the [579–594] peptide clearly confirm the presence of a disulfide bound within this sequence (Fig. 2D).

### Self-assembling and mechanical behavior

Reversible thermal responsiveness is a hallmark of native tropoelastin and many EDPs or ELRs [34, 35]. Here, we analyzed the thermal profile of a 1 mg/mL SEP solution using a dynamic light scattering approach (Fig. 3A). The data revealed that coacervation of SEP initiates above 30°C, which corresponds to the transition temperature (*T*_t_). A maximal coacervation was achieved at 60°C. The measured size of coacervates confirmed a temperature-dependent variation of the diameter of SEP aggregates as indicated in Table 1. The formation of larger aggregates in suspension above *T*_t_ caused an abrupt increase in turbidity. Indeed, aggregates below *T*_t_ (20°C) were of about 60 nm while they reached 13 µm above *T*_t_ (60°C). Upon cooling, the SEP recovered to a similar size as before heating, which proves the reversible transition capability of the molecule.

**Fig. 3.**
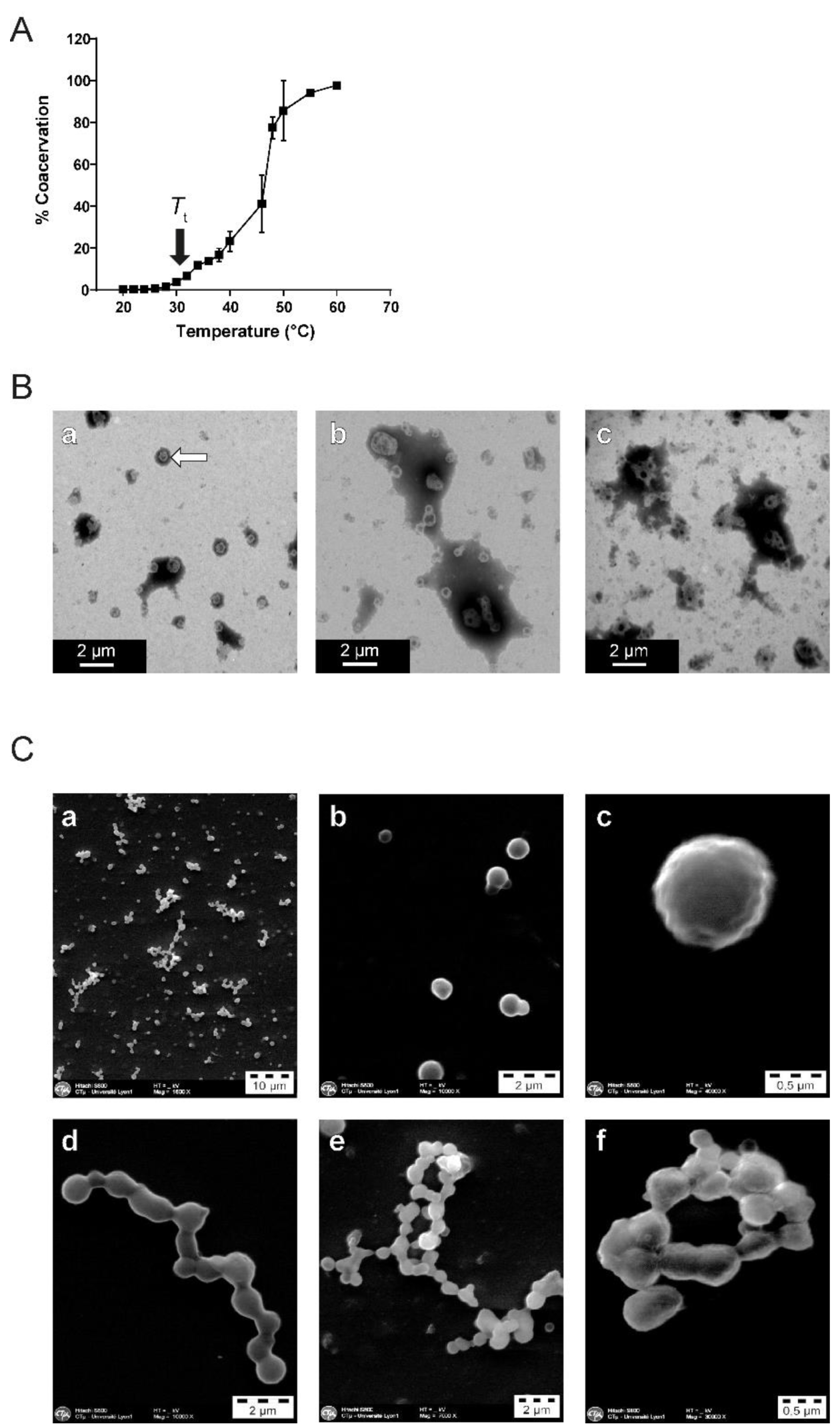
SEP self-assembles over *T*_t_. **A)** Coacervation profile of 1 mg/ml SEP solution in PBS (pH 7.4) in the temperature range 20- 60°C. Change in solution turbidity at each temperature expressed as a coacervation percentage of the overall maximum. **B)** Transmission electron microscopy imaging of SEP coacervates formed at 120 µg/mL (Ba), 240 µg/mL (Bb) and 2 mg/mL in PBS (Bc). **C)** Scanning electron microscopy images of SEP structures (Ca), monomers (Cb-c) and coalescent spheres (Cd-f)

**Table 1.**
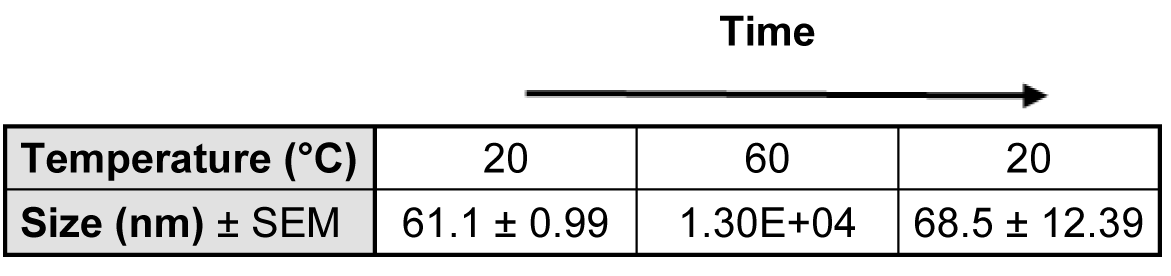
Coacervate size variations after heating and cooling a 1 mg/mL SEP solution.

|  |  |  |  |
| --- | --- | --- | --- |
|  | **Time** |  |  |
| **Temperature (°C)** | 20 | 60 | 20 |
| **Size (nm) ± SEM** | 61.1 ± 0.99 | 1.30E+04 | 68.5 ± 12.39 |

To further characterize the aggregation properties of SEP, 120 µg/mL to 2 mg/mL solutions were prepared in PBS and heated above *T*_t_ to induce coacervation. SEP was then visualized by transmission electron microscopy following negative staining (Fig. 3B). At 120 µg/mL, SEP formed small individual spheres of low internal electron-density (white arrow in Fig. 3Ba). At 240 µg/mL, spheres did not remain separate, but started to merge forming aggregates up to 2 µm in diameter (Fig. 3Bb). At 2 mg/mL, aggregates showed a higher external electron-density due to the merging of SEP individual spheres (Fig. 3Bc). Scanning electron microscopy indicated that the larger spheres observed at 2 mg/mL were due to merging (Fig. 3Ca). Edges of individual spheres were not regular (Fig. 3Cb-c). In addition to organize themselves in individual spheres, chimeric protein molecules coalesced together and connected like beads on a string (Fig. 3Cd), thus forming many varied geometrical shapes (Fig. 3Ce-f).

Since the thermal transition in solid state remains a reversible phenomenon, SEP aggregates were cross-linked to further address the mechanical behavior of the material at room temperature. Standardized tensile specimens were obtained by heating SEP solution over *T*_t_ in the presence of genipin inside silicone molds (Fig. 4Aa). The dark green samples displayed a regular dog-bone shape (Fig. 4Ab). However, the density of material was not homogeneous within the same sample due to the self-aggregation by sedimentation, showing areas of low density. These micro-defects could constitute points of brittleness and initiate material failure under stress, leading to an underestimation of the material’s strength (Fig. 4Ac, white arrows). Force-displacement curves were obtained by uniaxial tension until rupture. The elastic moduli (E) were computed by linear regression over the linear regions, i.e stretch ranging from 1 to 2 (Fig. 4B). Nominal rupture stress (Smax) and stretch (λmax) were extracted from the curves. Cross-linked SEP exhibited a linear mechanical behavior up to stretches of 2, with an average elastic modulus of 51.2 kPa and sample thicknesses varying from 245 µm to 437 µm (Table 2).

**Fig. 4.**
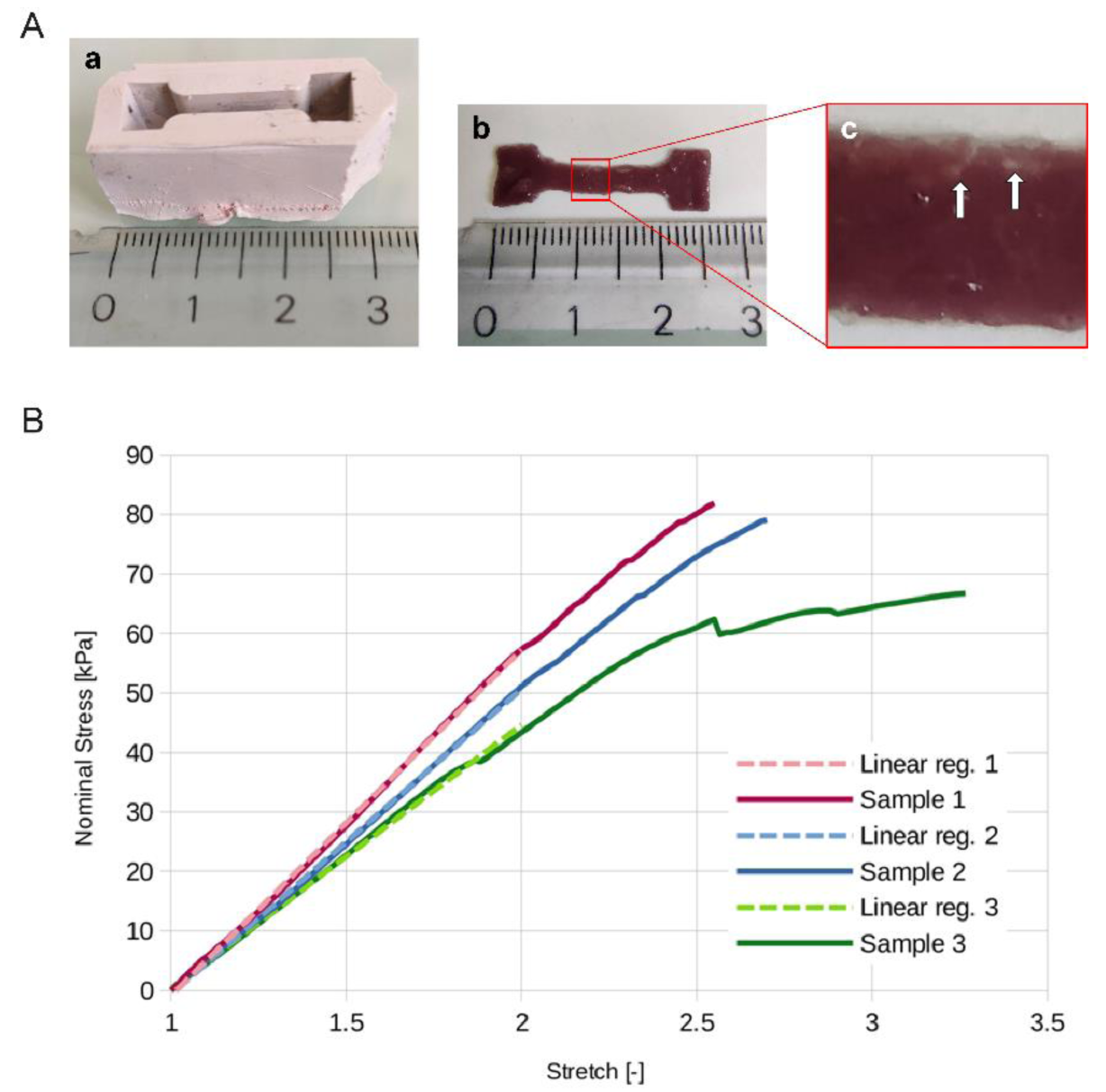
Cross-linked SEP has an elastic behavior. **A)** Cross- linked SEP specimen preparation for traction experiments. Silicone mold used (a). Result of SEP at 50 mg/mL supplemented with 50 µg/mL genipin, heated to 40°C overnight and then removed from the mold (b). Magnified image of an area showing the *density heterogeneity of a* sample (c). **B)** Stress-stretch curves up to rupture for the three specimens.

**Table 2.**
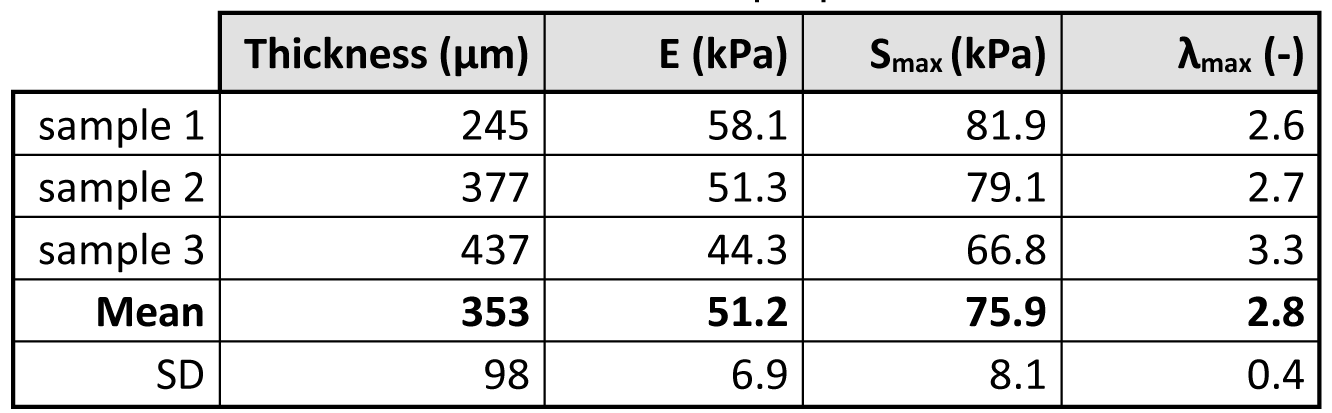
Thickness and mechanical properties from cross-linked SEP specimens.

| | Thickness ($\mu\text{m}$) | E (kPa) | $S_{\text{max}}$ (kPa) | $\lambda_{\text{max}}$ (-) |
| --- | --- | --- | --- | --- |
| sample 1 | 245 | 58.1 | 81.9 | 2.6 |
| sample 2 | 377 | 51.3 | 79.1 | 2.7 |
| sample 3 | 437 | 44.3 | 66.8 | 3.3 |
| **Mean** | **353** | **51.2** | **75.9** | **2.8** |
| SD | 98 | 6.9 | 8.1 | 0.4 |

### Biological activity of SEP towards vascular cells

ELRs are usually designed to incorporate biomaterials and are tested for their biocompatibility in their material-formulated form. Our objective is quite different here and aims at addressing a tropoelastin-like entity to reinforce elasticity naturally by the cells themselves. Therefore, we first analyzed intrinsic biological effects of SEP on vascular cells grown as monolayers.

Different concentrations of SEP, from 1 to 100 µg/mL, were introduced into the culture media of human umbilical vascular endothelial cells (HUVECs) and aortic smooth muscle cells (SMCs). Toxicity was first addressed by colorimetric metabolic assay after 48h (Fig. 5A-B). In these conditions, neither the HUVECs, nor the SMCs were affected by the treatment thus indicating no particular cytotoxic effect of the SEP. In the same way, the highest concentration was used to observe cell proliferation by monitoring cell density in culture over time (Fig. 5C- D). Again, SEP did not cause any significant change in the rate of proliferation of the two cell types. These results indicate that despite the strong representation of the VGVXPG hexa-motif in the SEP sequence, it does not appear to reproduce the proliferation-stimulating effects of free hexapeptides as observed in different cell types [36–38].

**Fig. 5.**
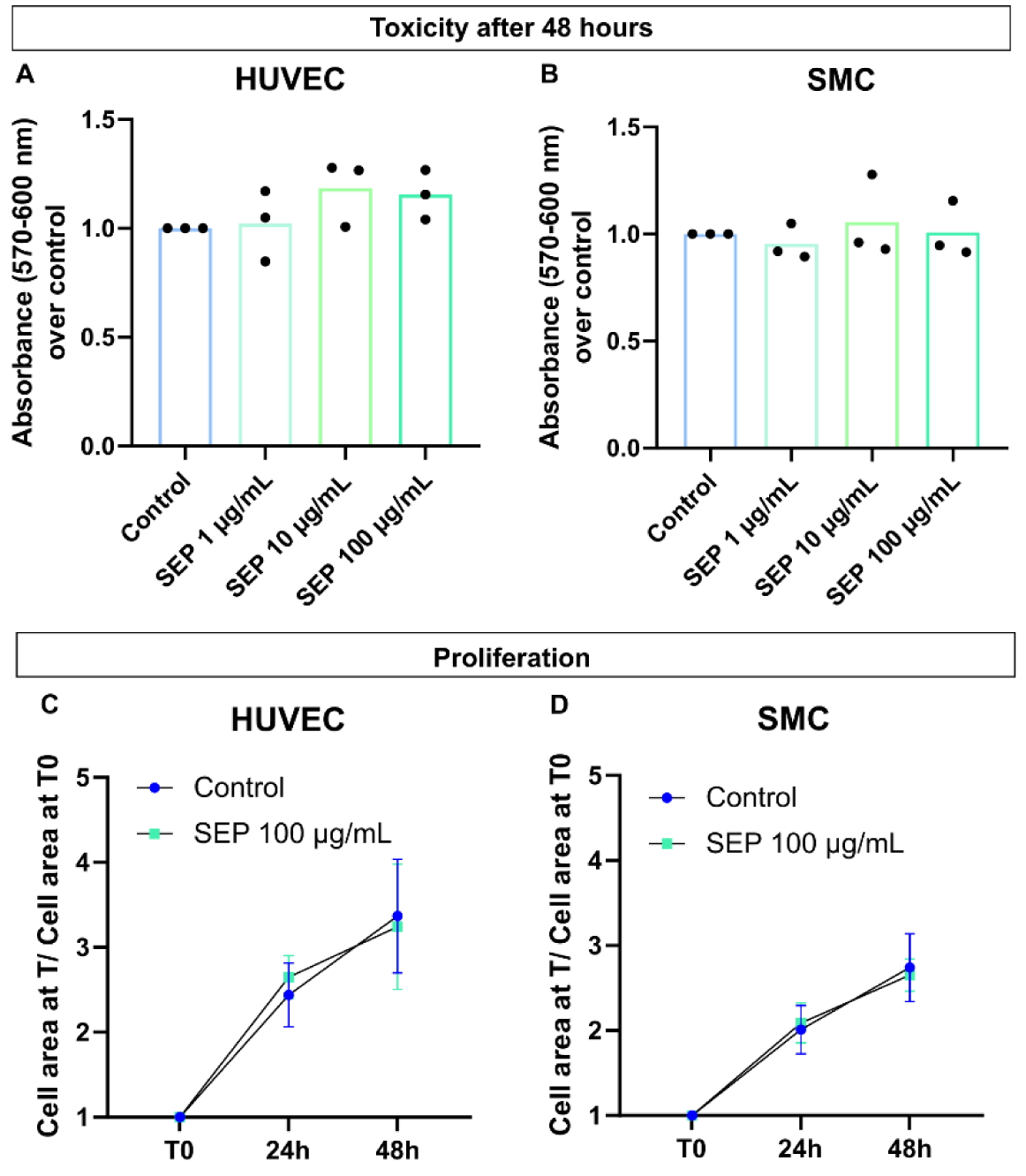
SEP is not toxic and does not impact vascular cell proliferation. **A-B)** Cell toxicity was assessed by adding 1-100 µg/ml of SEP into the culture medium of HUVECs **(A)** or SMCs **(B)** vascular cells and measuring their viability after a 48h exposure. 3 independent experiments were realized with 6 replicates of each condition. Black dots represent the mean of each independent experiment. **C-D)** Cell proliferation was also evaluated for 48h after the addition of SEP on HUVECs (**C**) and SMCs (**D**). The area occupied by cells at 24 or 48h was divided by the area occupied by cells at T0 to evaluate the progression of well occupancy in each condition. 3 independent experiments were performed with 3 replicates each. The mean of all three experiments is represented with error bars corresponding to standard deviation.

### Transendothelial passage and fiber formation

In order to reinforce vascular elasticity, SEP has to be delivered in the blood flow and integrate vessel walls. This concept requires that SEP is able to cross the endothelial barrier to be processed by SMCs and further incorporated into elastic lamellae. To address this, we used a co-culture model consisting in a cohesive monolayer of endothelial cells maintained in an insert above a culture of SMCs. Albumin coupled to a fluorescent dye (Fig. 6A) or SEP (Fig. 6B) were diluted in the culture medium above the endothelial cells. The integrity of the cellular junctions of endothelial cells was verified by the presence of vascular endothelial (VE)-cadherin at the level of the plasma membranes thus forming homogeneous honeycomb structures. A negative control consisting in 70 kDa fluorescent dextran was also used to check the barrier impermeability (not shown). The presence of fluorescent albumin (positive control) and SEP was sought at the level of the fibrillin-rich microfibrils produced by the underlying SMCs. While albumin is not found at the level of SMCs in the lower compartment, indicating that the protein did not passively adsorb, the SEP was immunodetected as fibrillar deposits at the level of fibrillin-1 structures. This result demonstrates that SEP is able to cross the endothelial barrier and to rearrange near the microfibrils formed by the SMCs by a mechanism that is probably more complex than adsorption.

**Fig. 6.**
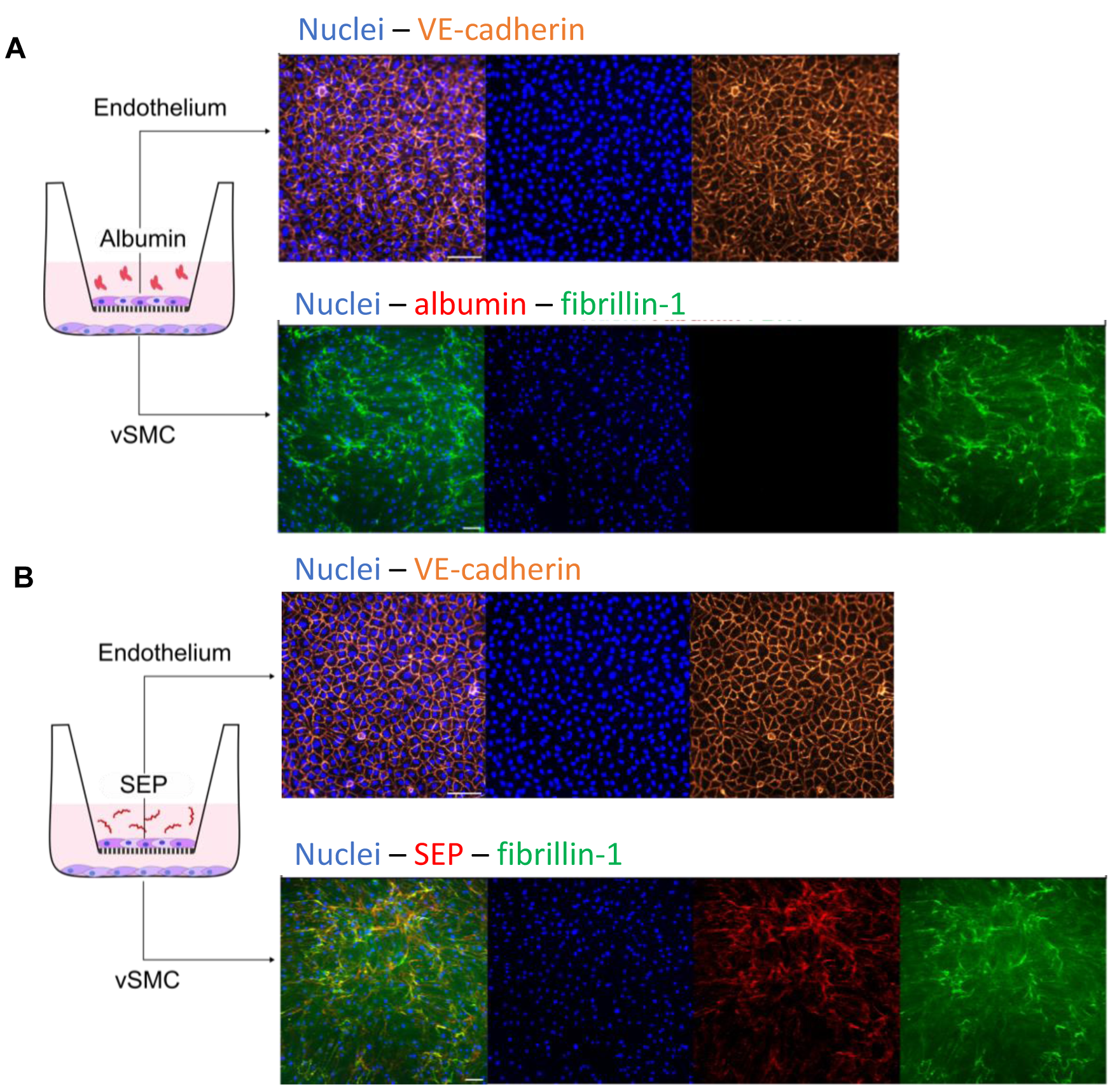
SEP is able to cross the endothelium and to integrate fibrillin-rich fibers in our *in vitro* co-culture model. We developed a co-culture system to assess the transendothelial passage of SEP, which consists in endothelial cells cultured on a porous membrane on an insert, that is then put on top of vSMCs cultured in a well, as illustrated on the left. Either fluorescence-coupled Albumin (250 µg/mL) **(A)** or SEP (250 µg/mL) **(B)** was added in the culture medium of the upper compartment. After 24h, both cell layers were analyzed. The integrity of the endothelial layer was confirmed through VE-Cadherin immunolabelling, and SEP passage through the structure reproducing the endothelial barrier and subsequent handling by the SMCs was revealed by fibrillin-rich fiber and Flag immunodetection. FBN1 = fibrillin-1; vSMC = vascular smooth muscle cells. Scale bar = 100 µm.

To better understand the management of SEP by SMCs, the protein was added in a single dose at 100 μg/mL into the culture medium of confluent SMCs. Its integration at the level of fibrillin-1 microfibrils was observed by anti-Flag immunofluorescence at 2, 8 and 15 days post-treatment (Fig. 7A-F). After 2 days, SEP partially co-localized with fibrillin-1 structures, and also formed smaller and more intense aggregates apparently aligned with finer fibrillin structures (Fig. 7D, white arrows). At 8 days, almost all the fibrillin structures were associated with SEP, and the intense small aggregates were still visible (Fig. 7E). At day 15, the intensity of the aggregates had decreased, and a large number of new fine structures of SEP-associated fibrillin covered all the SMCs (Fig. 7F). SEP could be associated to fibrillin structures either through specific passive interactions, or through active processes driven by SMCs. To determine whether the observed phenomenon was dependent on enzymatic activity, we used a LOX inhibitor (Fig. 7G-G’). In the presence of β-aminopropionitrile (β-APN), SEP aggregates were almost no longer visible, and only sparse co-localization with fibrillin structures could be observed. These results therefore demonstrate that SEP can be recognized by SMCs to be woven into microfibrils via the enzymatic activity of LOX. Interestingly, the fibrillar structures integrating the SEP seem stable since the SEP was still present and had not been discarded or digested by proteolysis 15 days after treatment. In this respect, however, SEP remains sensitive to degradation by main elastases such as MMP-12 and leukocyte elastase, but undoubtedly its highly repetitive sequence allows it to be quite more resistant than hTE is (Fig. S2).

**Fig. 7.**
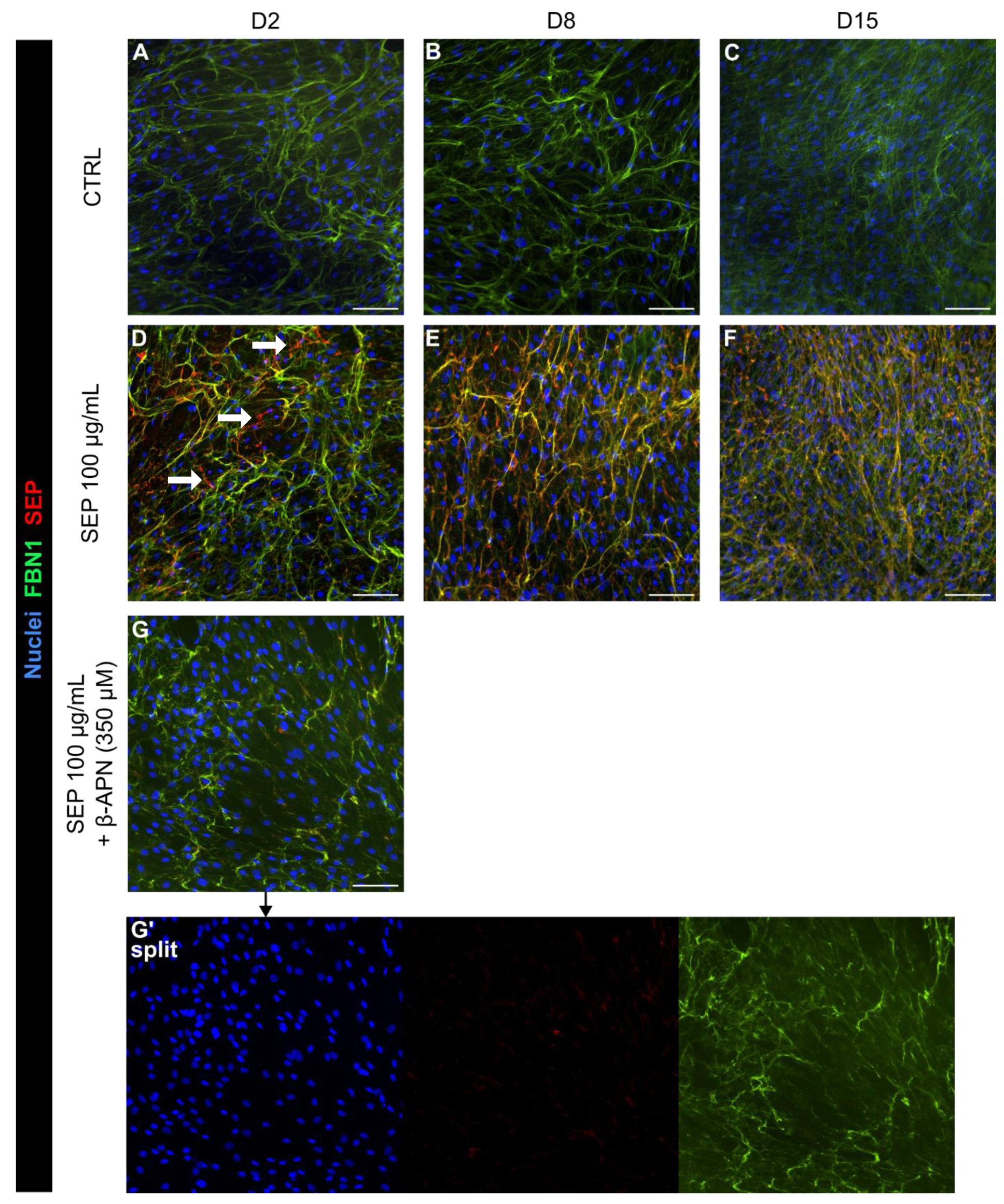
SEP integration into fibrillin-rich fibers persists over time and requires LOX activity. SEP was added into the culture medium of vascular smooth muscle cells for 2 days with or without β-APN (350 µM). Cells were then fixed at different time points, knowingly at day two (**A, D, G**), eight (**B, E**) and fifteen (**C, F**). Fibrillin-rich fibers were immunodetected in green and SEP in red. (**G’**) shows split channels of G, to highlight the decreased presence of SEP. FBN1 = fibrillin-1; β-APN = β-aminopropionitrile; LOX = lysyl oxidase. Scale bar = 100 µm.

### SEP behavior in the zebrafish model

The *in vitro* experiments suggest that SEP behaves as an analogue of hTE. To go further and initiate *in vivo* studies, we chose to use the zebrafish model that we recently reported as a relevant tool in the analyses of cardiovascular issues [39]. Specifically, we used the transgenic zebrafish reporter line Tg(*fli1a*:eGFP)/Casper, whose endothelial cells are fluorescent and enable vessel development observation in real time. In first intention, we evaluated the toxicity of SEP on dechorionated or non-dechorionated embryos (Fig. 8A). The embryos were incubated by immersion in increasing concentrations of SEP (25, 100 and 500 µg/mL) and their morphology and survival were monitored during 80h. A strong mortality was observed in presence of the chorion for the highest concentration of SEP. However, this is not related to a direct toxic effect of the protein on the physiology of individuals since no significant effect (development delay, pericardial edema, malformation of the tail) was observed using dechorionated individuals. We assume that it is more likely a physical phenomenon leading to accumulation of SEP in the chorion pores and preventing exchanges with the environment through this porous membrane. This assumption is notably reinforced by the presence of aggregates on the surface of the chorion likely to block the entry of the pore channels (Fig. 8A, left panel arrowheads) as previously described following immersion of zebrafish eggs in nanoparticle bath [40]. The safety of compounds derived from elastin is relatively established, however their inflammatory status is often questionable [41]. We therefore used the zebrafish reporter line Tg(*mpx:*eGFP)/Casper, whose neutrophils are labeled with GFP, to estimate the chemoattractant potential of SEP. SEP was injected into the yolk sac of 3 days post-fertilization individuals and neutrophil recruitment was monitored after 24 h (Fig. 8B). While many neutrophils were attracted to the injection site in the positive control using E. coli bacteria, no significant recruitment could be recorded for SEP in comparison with the vehicle injection (PBS). The inert inflammatory status of SEP was also confirmed in the human monocyte cell line THP-1 (Fig. S3). SEP or its degradation products, obtained by digestion with leukocyte elastase, did not induce an increase in the expression levels of pro-inflammatory cytokines (IL- 1β, IL-8 and TNFα) unlike the lipopolysaccharide (LPS) positive control. These results therefore argue in favor of an immunoquiescent nature of the SEP.

**Fig. 8.**
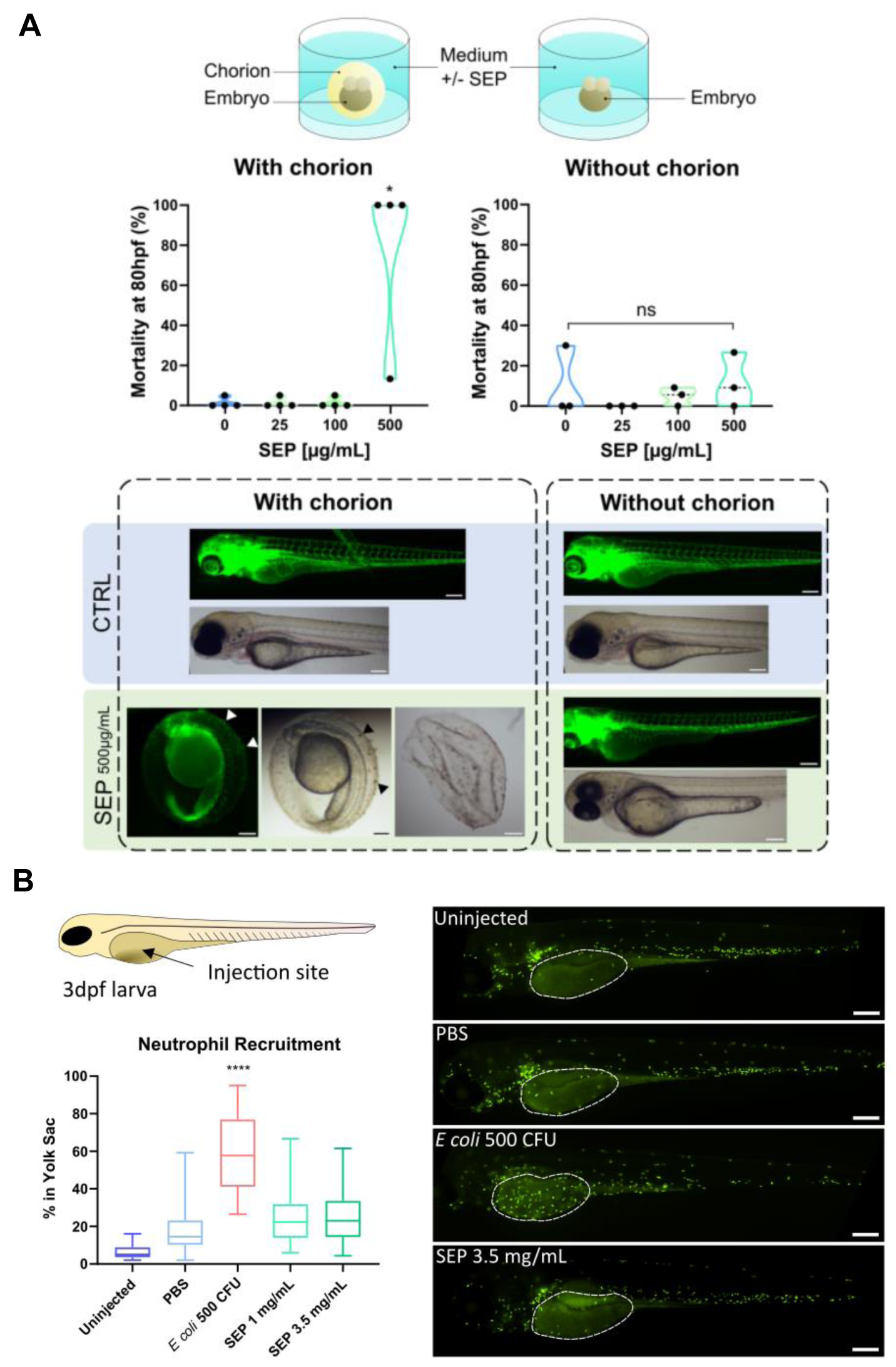
SEP does not have toxic effects on zebrafish when the chorion is removed and has no pro-inflammatory action. **A)** SEP toxicity was assessed *in vivo* by the Fish Embryo Toxicity Test. Zebrafish embryos were put in a culture plate containing E3 medium and SEP at increasing concentrations (here from 25 to 500 µg/mL). The effects were monitored for 80 hours. The chorion was either removed or not. Tg*(fli1a:*eGFP*)/*Casper zebrafish were used, to follow vascular development, as well as wild type AB/Tü zebrafish. Arrowheads point to SEP aggregates formed on the chorion. 4 (with chorion) and 3 (without chorion) independent experiments were performed with a total of at least 35 embryos per condition. Data were analyzed by the Kruskal-Wallis test; * = p<0.05, ns = non-significant. Scale bar = 200 µm. **B)** SEP pro-inflammatory potential was assessed by monitoring the recruitment of neutrophils 24h after injection into the yolk sac. PBS was used as a negative control and E. coli (500 CFU) as a positive control for recruitment. Tg(*mpx:*eGFP)/Casper zebrafish were used to follow neutrophils and the proportion of neutrophil recruited at the injection site (indicated by white dotted lines) was measured. 3 independent experiments were performed with at least a total of 25 fish per condition (except for the uninjected fish). Data were analyzed by the Kruskal-Wallis test; **** = p<0.0001 (compared to PBS injection). dpf = days post fertilization; CFU = colony forming unit; Scale bar = 200 µm.

Finally, SEP coupled with a fluorescent dye was microinjected directly into the blood flow of the Tg(*fli1a:*eGFP)/Casper zebrafish strain to observe its distribution in the vasculature for 96h (Fig. 9). As early as 3 h post-injection (hpi), SEP was found in all visible vessels, including the caudal vein, the dorsal aorta and the intersegmental vessels. More particularly, we noted the presence of dots which could correspond to the aggregation of SEP along the vessels. At the same time, Dextran (70 kDa), used as a negative control, also showed staining throughout the vascular tree, but no focal dot could be observed, giving way to diffuse staining in the vessels. After 24 h, SEP aggregates became more defined and seemed to accumulate in the caudal vein of the individuals, where the vascular tree is the most tortuous. At this same time, the labeling of Dextran appeared diffuse and decreased in intensity compared to the time-point at 3 hpi, which may result from a clearance of the compound. Further analysis by confocal microscopy clearly highlighted the incorporation of SEP inside the vessel and beyond the endothelial barrier, thus ruling out the passive deposition of SEP in tortuous area (Fig. S4). At 96 hpi, SEP aggregates remained visible but of less intensity, whereas the Dextran was no longer detected at all. The decrease in intensity of the SEP aggregates could come either from a degradation of the exogenous material, or from the release of the fluorochrome following SEP incorporation into the blood vessel, or from a bleaching due to the oxidation of the fluorescent group over time. In conclusion, these *in vivo* results corroborate the observations made previously in *in vitro* models where SEP accumulates and persists over time. This phenomenon therefore suggests that the protein would be processed to be integrated into the extracellular matrix of the vessels, although additional analyzes will be necessary to confirm this.

**Fig. 9.**
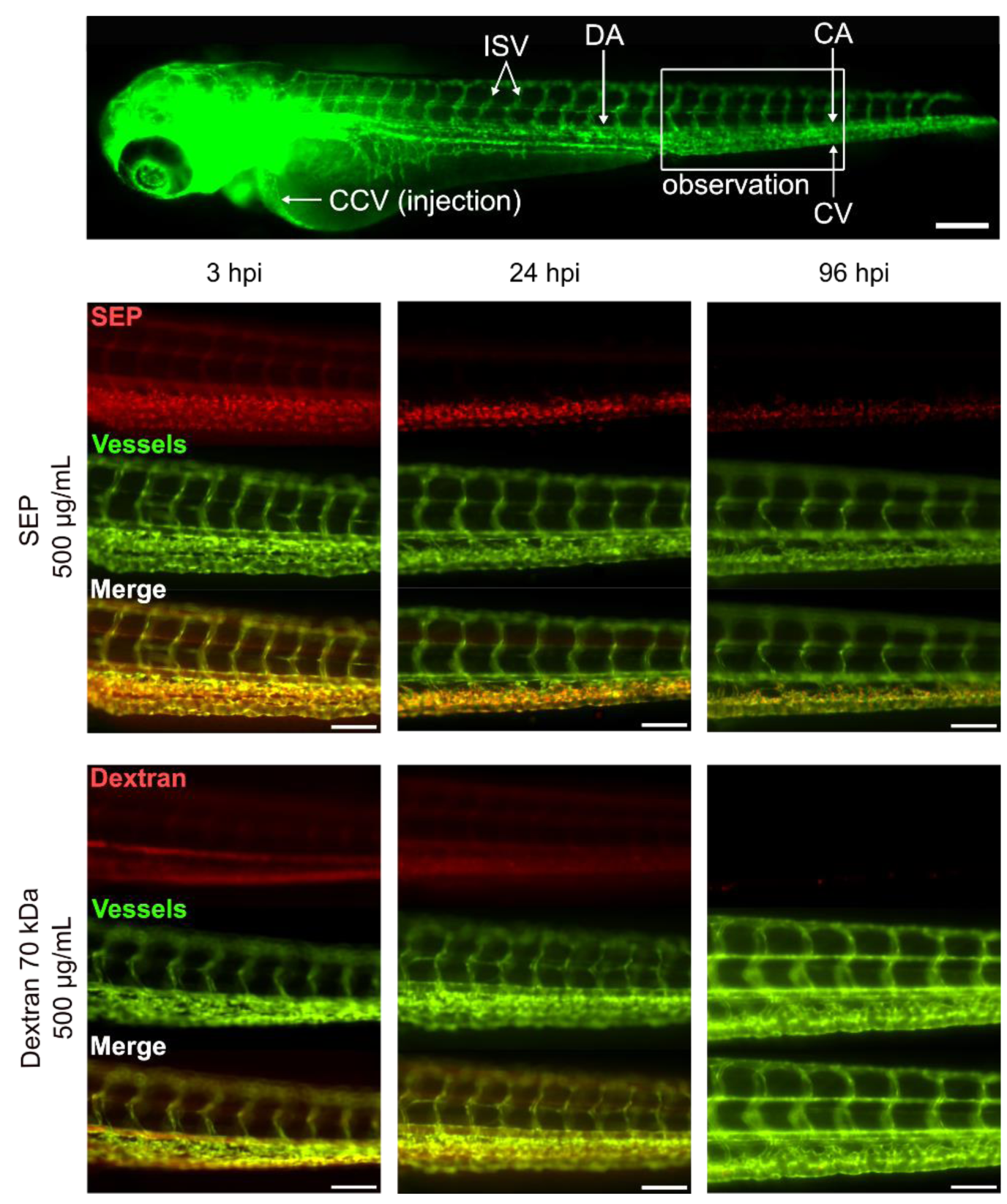
SEP is retained in vessels after intravascular injection in zebrafish embryos for at least 96h. Zebrafish embryos Tg(*fli1a*:eGFP)/Casper were injected with either SEP or Dextran 70 kDa (control) at 2 days post fertilization and monitored by epifluorescence microscopy for 4 days to follow the injected molecules. Vessels appear in green and injected molecules are coupled with red fluorophores (SEP-Rhodamine and Dextran-Rhodamine). The injection was performed in the CCV and the observation zone corresponds to the tail of the embryo which is delineated with a white rectangle. hpi: hours post injection; CCV: common cardinal vein or duct of Cuvier; DA: dorsal aorta; CV: caudal vein; CA: caudal artery; ISV: intersegmental vein. Scale bar = 100 µm.

## Discussion

The objective of this study was to characterize a polypeptide inspired by the human tropoelastin and to evaluate its ability to associate with elastic fibers for vascular repair purpose. The repair of damaged elastic fibers by adding tropoelastin had already been proposed following the development of a gene coding for a soluble recombinant common form of tropoelastin expressed in E. coli [42]. This recombinant tropoelastin had then been optimized and its incorporation into endogenous elastic fibers had been demonstrated in cultures of rat fibroblasts [43]. If this construction and others have largely contributed to the understanding of the molecular mechanisms related to the complex assembly of elastic fibers, to our knowledge however, no therapeutic application has been proposed to date. Several parameters can explain this translational lock. First concerning the intrinsic properties of hTE itself, such as the sensitivity of hTE to proteolysis and subsequent effects of the derived fragments. The multiple isoforms that should be delivered specifically to each tissue to recover the initial properties of the elastic fibers is also an almost impossible objective to achieve. Practical aspects must be taken into consideration, such as its production and purification often based on reverse phase chromatography using solvents that are not very compatible with pharmaceutical development or requiring very heavy quality controls, although the production price has been estimated as low as US$0.70 / mg under good manufacturing practice conditions [44], this does not include pharmaceutical qualification. Finally, the possibility of delivering the protein to the target connective tissues remains a major challenge. For these reasons, hTE has in fact found another very important sector of applications in the formulation of biomaterials which fall under the slightly less restrictive regulatory issues of medical devices. We can report, among other great successes in murine models, the functionalization of vascular stents [45], or the grafting of bioresorbable vessels [46], where hTE prevents restenosis and hyperplasia. However, to our knowledge, these devices have never been validated for human applications.

The SEP used in our study takes up few conformational characteristics of hTE, in particular the alternation of hydrophobic and cross-linking domains. As retrieved in many other hTE- derived polypeptides, the hydrophobic domains consist of aliphatic amino acids (valine-glycine-leucine-alanine) and a proline repeated at regular intervals, here every 6 residues. The space between the proline residues is essential for the reversibility of the coacervation and a period of 6 residues, as found in domain 24 of hTE, allows an optimal reversibility of 98% [47]. From 9 residues and beyond, as found in domain 30 of hTE, coacervation results in the formation of stable amyloid-β structures [48, 49]. These rigid structures are therefore unwanted and can contribute to the development of neurological pathologies such as Alzheimer’s disease [50]. Our NMR results, which were recorded from SEP coacervates, indicate that even in the solid state, SEP retains a significant degree of flexibility at the proline residue level. These data are in good agreement with the recent characterization of elastin in solid-state NMR coupled with Fourier Transform Infrared experiments reporting that the flexibility and elasticity of native elastin are linked to the *cis-trans* isomerization of the Xaa-Pro imide bonds [51]. Between the hydrophobic domains, we interspersed alanine-lysine-rich cross-linking domains corresponding to a motif found in domain 19 of hTE and conserved across many species. To allow cross-linking, these domains must have an α-helix or polyproline II (“extended helix”) conformation, thus placing two lysines sufficiently close to cause the formation of desmosine and isodesmosine cross-links [52]. Circular dichroism and NMR analyzes indicate that SEP possesses such structures and would therefore be prone to cross-linking by endogenous machinery.

To determine the mechanical properties of SEP, we therefore used genipin, a natural non-toxic agent having already been shown to be effective in the crosslinking of lysines of elastin and its derivatives at the level of the crosslinking domains [53]. Cross-linking has the effect of insolubilizing the material, thus making coacervation irreversible, as in the process of maturation of elastic fibers. The cross-linked SEP samples could therefore be mechanically analyzed by a tensile test demonstrating a wide range of linear elongation and an elastic modulus around 50 kPa. However, the values of elastic modulus and rupture strength cannot be compared with other studies in which the samples were centrifuged in a highly saline medium (1.5M NaCl) to force coacervation and densify the material, thus offering much higher elastic modulus values of the order of 1MPa [54, 55]. We opted for concomitant cross-linking with sedimentation of the coacervates in a physiological buffer to bring us closer to the native conditions for the elastic fiber assembly, nevertheless this process offers a less dense and less homogeneous material with areas of fragility. Though, beyond the absolute elastic modulus values, we will mainly retain the linear elastic behavior of the SEP-based material reflecting the global SEP mechanics.

We then showed that SEP added to the culture medium of human vascular cells did not impact the survival or proliferation *in vitro*. At the endothelial cell level, the reported effects of hTE and certain derived peptides in the soluble phase mainly concern the release of NO with consequences on vasorelaxation [56, 57]. Recently, a positive effect of hTE on HUVEC tubulogenesis has been reported [58]. In this study, the authors showed that this effect was indirect and responded to a signal provided by the mesenchymal stem cells treated with hTE. Regarding SMC, we could have expected a decrease in cell proliferation. Indeed, SMCs deficient in elastin, as in the case of WBS or SVAS, have a low level of tropoelastin synthesis associated with an increase in their proliferation, this phenotype was reversed by adding hTE to their culture medium [17]. However, this effect was observed for concentration of 1 mg/ml, 10 to 100-fold higher than what we tested for SEP. The most interesting point of our results on vascular cells certainly concerns the ability of SEP to cross the endothelial barrier and further be incorporated into fibrillar deposits by SMCs in a co-culture model. While it is known that elastin-derived peptides are able to cross anatomical barriers to even be found beyond the blood-brain barrier to contribute to Alzheimer’s disease as mentioned above [50], we did not find any element reporting such a mechanism for hTE. The mechanism by which SEP crosses the endothelial layer is not elucidated but we found SEP inside HUVECs following treatment, suggesting a mechanism of transcytosis (not shown). Finally, SEP reaching the SMCs is processed and deposited at the level of fibrilin-1-rich fibrillar structures. Kozel and coworkers had described that hTE could passively associate with a native matrix in the absence of living cells via sulfated proteoglycans [59]. This constitutes the early phase of tropoelastin deposition, however, in the presence of an inhibitor of LOX activity (β-APN), no deposition of SEP aggregates could be observed. This result therefore clearly demonstrates an active cross- linking of the SEP at the level of the SMC matrix, rather than a passive adsorption of the protein onto pre-existing fibres. Also, this phenomenon evolves favorably over time even without further addition of SEP, definitely demonstrating an active incorporation. Although observations at the ultrastructural scale would be necessary to confirm this, SEP seems to be recognized by SMCs in a way similar to tropoelastin and correctly addressed at the level of the extracellular matrix.

In the last part of our study, we applied SEP in the zebrafish model to observe its behavior *in vivo*. Our results show no toxicity of SEP even for high doses up to 500 µg/mL in the medium of dechorionated embryos in compliance with the guidelines n°236 proposed by the OECD for this test. Zebrafish is considered as a good, inexpensive and highly scalable model for *in vivo* drug testing in a preclinical trial and predicting the toxicity of therapeutic molecules in humans [60–62]. Besides the number of descendants generated from a single spawn (100-200 eggs per female and per week) and the small size of the embryo which enables to perform high throughput experiments without to use high amount of drug for testing, small molecules can be actively absorbed from the water by the embryo which facilitates drug administration. Also, numerous studies exist demonstrating that zebrafish is sometimes a better model, for predicting the toxic effects of drugs on humans compared to some mammalian models [63–65]. Interestingly, no defect, including pericardial edema, intersegmental vessel malformation or blood flow slowing, was observed following embryo immersion in SEP solution which suggests the absence of adverse effect of SEP on zebrafish cardiovascular system [66]. Following the evaluation of SEP toxicity in zebrafish embryos by immersion, we also injected the SEP into the duct of Cuvier - which is the largest vessel of this animal model - of 2 dpf embryos, and monitored the biodistribution of this molecule for 96h. This delivery method was chosen to come as close as possible to a drug delivery system capable of repairing the wall of vessels lacking elasticity.

In contrary to the Dextran used as control, SEP was accumulating into the caudal vein connected to the posterior cardinal vein, which collects all the blood from the body and transport it back to the heart. This vasculature forms an interconnecting network of venous tubes, in which the vascular topology results in locally high shear gradients and slower blood flow compared to the dorsal aorta [67]. This could favor the interaction between molecules in the bloodstream (and thus the SEP) and the endothelial cells. This particular tortuous topology found in zebrafish embryo vasculature could be compared to abnormal vasculature found in aortic aneurysm, supravalvular aortic stenosis and in aging arterial wall [68]. Therefore, this last result implies the SEP deposition in pathological conditions and reinforce the interest of the zebrafish model to study drug for vascular pathologies.

With the fantastic emergence of genome editing over the last 20 years, strategies aiming at reprogramming cells for the proper production of proteins, and particularly Crispr/cas9 technology, has become a hot topic [69, 70]. However, this strategy might not be relevant for elastin deficiency since elastin is a particularly stable protein and for which time-limited administration/activation of production could be sufficient for the improvement of mechanical defects. Recently, the intradermal delivery of mRNA encoding for tropoelastin, in solution with lipofectant, has shown significant increase in elastin levels in pig skin *ex vivo*, but again this involves transfection of specific cells in the body which is difficult to consider on arterial SMCs which are buried deep in the vessel [71]. In this respect, a strategy using a protein capable of crossing the endothelial barrier and binding directly to the fibers in the tunica media in situ therefore remains relevant. Recently we reported the use of SEP as a single intravenous injection in an elastin haploinsufficient mouse model [72]. Aortic physio-mechanical improvements were shown for high blood pressure values at 3 days post-injection, but the presence within the vascular walls could not be observed. We had concluded that it could be a biological rather than a mechanical action of the SEP. Considering the results brought by the present study, SEP could nevertheless reach the vascular wall as shown in zebrafish, but probably in lower proportions in mice. It would be highly appropriate to perform multiple injections to increase SEP incorporation into the vascular walls until detectable levels to reach more significant mechanical input, even at low blood pressure.

## Conclusions

In summary, the SEP is a novel polypeptide that mimics tropoelastin and recapitulates the physicochemical, mechanical and biological behavior of the native protein without inducing cytotoxicity or systemic inflammation. It can be administered intravenously and actively

crosses the endothelial barrier to reach vascular smooth muscle cells and incorporates into the extracellular matrix. These results encourage us to further validate its potential in relevant mammalian models to improve arterial mechanical parameters when these are compromised.

## Experimental procedures

### Production and purification implementation

#### SEP production

Gene synthesis, encoding for SEP, was outsourced to GenScript (Piscataway, USA). The DNA sequence (1794 pb) was optimized for an E. coli prokaryotic system expression. The gene was cloned into the pET30a prokaryotic expression vector (Promega, Charbonnières-les-Bains, France) between *NdeI* and *SalI* sites.

Recombinant vector digestion with *EcoRV* restriction enzyme enabled to validate the purification of the right plasmid, prior sequencing (Eurofins Genomics France, Villeurbanne, France). pET30a-SEP vector was stored at -20°C prior to transform BL21 (DE3) competent bacteria (New England Biolabs, Évry-Courcouronnes, France).

The pET30a-SEP recombinant vector was used to transform BL21 (DE3) competent E. coli bacteria selected on Luria Bertani (LB) both-agar plates containing 50 µg/mL kanamycin (Sigma–Aldrich, Saint-Quentin-Fallavier, France). A single colony was used to inoculate 5 mL LB containing 50 µg/mL kanamycin overnight at 37°C under constant stirring. The culture was diluted 1:100 and SEP expression was induced at the exponential growth phase by the addition of 1 mM isopropyl-β-D-thiogalactopyranoside (IPTG) (Sigma–Aldrich) for 4h at 30°C under constant stirring. Alternatively, for large batch production, the preculture was inoculated at 1% (v/v) in 2 L of LB-based auto induction medium (Formedium, Hunstanton, UK) supplemented with 50 µg/mL kanamycin, and cultured overnight at 37°C under constant stirring. Similarly, for NMR experiments, ^13^C-^15^N uniformly labeled SEP was prepared in 2 L of minimal M9 medium containing 2 g/L of ^13^C-glucose and ^15^NH4Cl.

Bacteria were then harvested by centrifugation 4,000 g/4°C/10 min and the resulting cell pellet frozen overnight at -80°C. Bacteria were resuspended in water and subjected to mechanical cell disruption (Cell disruptor TS2, Constant Systems, Daventry, UK). Soluble and insoluble fractions were harvested by centrifugation 15,000 g/4°C/20 min and analyzed by Coomassie staining of sodium dodecyl sulfate–polyacrylamide gel electrophoresis (SDS-PAGE).

#### SEP purification

SEP was purified from the soluble E. coli lysate by inverse transition cycling (ITC) method, as previously reported by Chilkoti and Meyer [32]. Briefly, lysate was buffered with 50 mM Tris-HCl pH 8.8, and 0.2% (w/v) polyethylenimine (Sigma-Aldrich) was added under agitation for 30 min at 4°C. The precipitate was discarded by centrifugation 15,000 g/4°C/10 min. The supernatant was warmed to 40°C for 10 min with 1 M NaCl prior centrifugation at 15,000 g/40°C/5 min. The supernatant was discarded, and the pellet containing SEP was resuspended in cold phosphate-buffered saline (PBS) at pH 7.4 containing 0.5 M NaCl, and incubated on ice for 10 min to end the first ITC. A total of three rounds of ITC were applied. The final pellet was resuspended in 0.9% (w/v) NaCl or PBS. A quality control was conducted by silver staining of SDS-PAGE. The presence of endotoxin was quantified by limulus amebocyte lysate testing (Endosafe, Charles River, Ecully, France) and osmolarity was measured by osmometer (OsmoPRO, Advanced Instruments, Paris, France). The purified SEP was passed through a 0.2 µm pore size syringe filter for sterilization and quantified using the BCA Protein Assay kit (Thermo Fisher Scientific, Les Ulis, France). Aqueous solutions were stored at -20°C until further use.

#### Western blotting

1 µg of SEP was subjected to SDS-PAGE. Proteins were transferred onto poly(vinylidene fluoride) membrane using Trans-Blot Turbo Transfer System (Bio-Rad, Marnes-la-Coquette, France). After transfer, the membrane was blocked 1 h at room temperature with 5% (w/v) non-fatty milk in tris-buffered saline with Tween-20 at 0.1% (v/v) (TBS-T). Membrane was further incubated with anti-FLAG (1/5000, F1804, Sigma-Aldrich) in 2% (w/v) non-fatty milk in TBS-T overnight at 4°C. The membrane was rinsed three times with TBS-T and then incubated with goat anti-mouse IgG coupled to horse radish peroxidase (1/10,000, 1706516, Bio-Rad) in TBS-T for 1 h at room temperature. The membrane was rinsed five times in TBS-T and revealed with enhanced chemiluminescent substrate (Super Signal West Pico, Thermo Fisher Scientific). Images were acquired with FUSION FX imager (Vilber-Lourmat, Collégien, France).

### Structure characterization

#### Circular Dichroïsm

Far-UV CD spectra of SEP were recorded with a Chirascan spectrometer (Applied Photophysics, Leatherhead, United Kingdom) calibrated with 1*S*-(+)-10-camphorsulfonic acid. Measurements were carried out 20°C to 60°C in a 0.1-cm-path-length quartz cuvette (Hellma, Müllheim, Germany), with a SEP concentration of 0.2 mg/mL in PBS at pH 7.4. Spectra were measured in a 180 nm to 260 nm wavelength range. Spectra were processed, baseline corrected, smoothed, and converted with Chirascan software. Spectral units were expressed as the mean molar ellipticity per residue.

#### Nuclear Magnetic Resonance

^13^C-^15^N SEP was dialyzed in H2O after purification and lyophilized. 66 mg of protein were resuspended in 860 µL of PBS buffer and placed in a bath at 40 °C for 15 min for coacervation. The coacervated sample was filled into a thin-wall 3.2 mm rotor by ultracentrifugation (1 h at 100,000 *g*, 40 °C) using an NMR filling tool. Before closing the rotor, 1.5 µl of DSS were added for chemical shift referencing.

NMR experiments were recorded on a 800 MHz spectrometer at a 17.5 kHz magic-angle spinning (MAS) spinning frequency and at temperatures of 4 °C and 40 °C. The temperature difference did not affect the NMR spectra. 2D ^13^C-^13^C DARR spectrum was recorded with a mixing time of 40 ms and 8 scans for 15h45. 2D ^1^H-^13^C-INEPT spectra was recorded with 16 scans for 8 h 06 min. 2D ^13^C-^13^C INEPT-TOBSY [73] was recorded with 32 scans for 15h45 at 15 kHz MAS rate and a TOBSY mixing time of 6.2 ms.

#### Mass Spectrometry

After alkylation with 10 mM iodoacetamide for 45 min at room temperature, SEP was digested overnight at 37°C with sequencing grade pepsin (Promega) at a 1/20 (pepsin/SEP) mass ratio. Peptide sequences were determined by mass spectrometry performed on a LTQ Velos ETD instrument (Dual Pressure Linear Ion Trap, ETD activation mode) equipped with a nanospray source (Thermo Fisher Scientific) and coupled to a U3000 nanoLC system (Thermo Fisher Scientific). A MS survey scan was acquired over the m/z range 300-1600 in Zoom scan resolution mode. The MS/MS scans were acquired in Normal resolution mode over a dynamic m/z range for the 10 most intense MS ions bearing a charge equal or higher than 2. Acquisition parameters for the ETD MS/MS spectra were set as followed (i) reaction time to 100 ms, (ii) isolation width of the precursor ion (z=+3) to 2 amu. The spectra were recorded using dynamic exclusion of previously analyzed ions for 0.5 min with 50 millimass units (mmu) of mass tolerance. The peptide separation was obtained on a C18 PepMapmicro-precolumn (5 μm; 100 Å; 300 μm x 5 mm; Dionex) and a C18 PepMap nanocolumn (3 μm; 100 Å; 75 μm x 200 mm; Dionex) using a linear 60 min gradient from 0 to 40% of solvent B, where solvent A was 0.1% HCOOH in H2O/CH3CN (95/5) and solvent B was 0.1% HCOOH in H2O/CH3CN (20/80) at 300 nL/min flow rate.

Protein identification was performed using the SEQUEST Algorithm through the Proteome Discoverer software v1.4 (Thermo Fisher Scientific) against SEP sequence. The ETD MS/MS spectrum was validated manually.

### Self-assembling

#### Aggregate size measurement by Dynamic Light Scattering

SEP coacervation and aggregate size in solution was measured using a Malvern-dynamic light scattering Zetasizer Nano S ZEN1600 (Malvern Panalytical, Vénissieux, France), for a range of temperatures between 20 and 60°C, with a stabilization time of 5 min. SEP sample were prepared at 1 mg/mL in PBS, pH 7.4. 11 runs were performed for each sample to determine the particle aggregate size, in order to obtain a final average value at constant temperature.

#### Electron microscopy

For transmission electron microscopy, SEP was dissolved at different concentrations (120 µg/mL to 2 mg/mL) in cold PBS (4°C). All solutions were equilibrated at 42°C for 15 min to generate coacervation. A formvar-carbon film grid (Electron Microscopy Sciences, Hatfield, United Kingdom) was deposited on a droplet of each sample at 42°C. After 2 min, excess solutions on the grid was blotted and the adsorbed samples fixed with a 2% glutaraldehyde solution, for 2 min at 42°C. Samples were washed three times with distilled water, stained for 10 sec with 2% phosphor tungstic acid pH 7 (1M KOH) and air-dried. Observations were performed with a Philips CM120 transmission electron microscope, equipped with a digital CDD camera (Gatan), at the Centre Technique des Microstructures, University of Lyon 1.

For scanning electron microscopy, all solutions were equilibrated at 42°C for 15 min to facilitate coacervation, spread on glass coverslips (Braunschweig, Germany), and air-dried at 42°C for 15 min. The resulting coverslips were subsequently repeatedly washed in PBS prior dehydration in graded ethanol solutions (50% to 100%). All samples were mounted on aluminum stubs, followed by thin (∼3 nm) copper sputtering (Bal-tec MED 020, Leica Microsystemes SAS, Nanterre, France). Imaging was carried out using a Hitachi S800 FEG, scanning electron microscope.

### Characterization of mechanical properties

#### Genipin cross-linked SEP stretch specimens

Purified SEP was dissolved in PBS (150 mM NaCl, pH 7.4) to a final concentration of 50 mg/mL at 4°C. Samples were coacervated at 40°C in silicone molds in presence of 50 mM aqueous genipin (Sigma-Aldrich) and cross-linked overnight at 40°C. Samples were removed from the mold and preserved in PBS, pH 7.4, containing 20 % EtOH.

#### Uniaxial tension tests

Uniaxial tension tests until rupture were performed in PBS at room temperature on an Instron 3343 tension machine (Instron, Élancourt, France) at 10 mm/min, fitted with a Futek LSB210 100g load cell (Futek, Irvine, CA, USA). Nominal stress was computed as the ratio of the force and the initial cross-sectional area of the sample, using the dimensions of the width of the mold and thickness measured using optical interferometry (Table 2). Stretch was computed from the ratio of the initial length of the sample and its current length, determined by the jaw displacements.

### Cell Culture and *in vitro* assays

Human Umbilical Vein Endothelial Cells (HUVEC; CC-2519) and human Aortic Smooth Muscle Cells (AoSMC, CC-2571) purchased from Lonza were cultured in EGM (CC-3124) and SmGM- 2 media (CC-3182), respectively (Lonza, Basel, Switzerland).

#### Cell survival

Cell viability was evaluated using PrestoBlue reagent (A13261, Thermo Fisher Scientific). SEP was added at T = 0h and the medium was not changed before the test performed at T = 48 hours. At this time, the medium was removed and cells were incubated for 1 hour with diluted PrestoBlue, as recommended by the provider guidelines. The signal was then measured by absorbance using a microplate reader (AMR-100, Allsheng) at 570 and 600 nm, in new plates. Results were obtained by subtracting blank to all data, subtracting the 600 nm absorbance from the 570 nm, and then normalizing everything over the mean of control wells. 6 replicate wells were attributed to each tested condition. The experiment was repeated 3 times.

#### Cell proliferation

SEP was added at T = 0h and the medium was not changed during the experiment. Cell proliferation was evaluated using the Cytonote-6W technology (Iprasense, Clapiers, France). Images of cells were taken at 24 and 48 hours and proliferation was assessed by evaluating the area expansion of cultured cells over time, using the ImageJ software. 3 replicate wells were attributed to each condition tested. The experiment was repeated 3 times.

#### Co-culture in vitro model

HUVECs were cultured on an insert device with a porous membrane (0.4 µm – CLS3801; Merck, Darmstadt, Germany), seeded directly at confluency with complete EGM medium up and down for 2 days. SMCs were cultured for 3 to 5 days in 6 well plates in parallel, with 2 to 3 Softslips (coverslip with 4 kPa hydrogels coated with collagen; SS24-COL-4-PK; Cell Guidance Systems, Cambridge, UK) distributed in each well. At day -1, the HUVEC inserts were placed upon the SMCs in culture with EGM medium in the upper compartment and EBM (EGM without complement) in the lower compartment. These culture parameters were selected to obtain a permeability of the endothelial layer for which 70 kDa Dextran cannot cross. At day 0, the passage test was performed. SEP or fluorescently labeled Albumin (FITC-Albumin, A23015, Thermo Fisher Scientific) was added in the upper compartment at 250 µg/mL in 500 µL EBM medium. 24 h later, both the endothelium layer on the insert and the SMCs on the Softslips were collected to perform immunofluorescence. (To be noted: IF image colors were inverted so that all fibrillin-fibers appear in green and Albumin/SEP in red, so that FITC-Albumin is in fact in red).

#### SEP integration into fibers

SMC were cultivated on 4 kPa Softslips (same as above) with SmGM until confluency and then in SmBM (SmGM without complement) + 2% (v/v) FBS and antibiotics from the Bulletkit for SmGM (CC-4149, Lonza, Basel, Switzerland). SEP was added once at the beginning of the experiment at 100 µg/mL. For the β-APN condition (A3134, Sigma-Aldrich), cells were treated with 350 µM β-APN one hour before adding SEP. The treatment was renewed 24 h later by adding 10µL of a 100X solution in the medium containing SEP (1 mL). Softslips were collected at day 2, 8 et 15 to perform immunofluorescence.

#### Immunofluorescence

Cells were fixed in 4% (w/v) paraformaldehyde in PBS at room temperature for 15 min and rinsed 3 times with PBS, then permeabilized with 0.1% (v/v) Triton X-100 for 10 minutes on agitation and rinsed 3 times, then incubated with 5% (w/v) BSA for 45 minutes on agitation. The following antibodies were used: anti-VE-Cadherin (1/500, 55566, BD Pharmingen, Le Pont de Claix, France), anti-FLAG (1/500, F1804, Sigma-Aldrich); anti-FBN1 (1/500, HPA021057, Sigma-Aldrich). All primary antibodies were incubated with 0.5% (w/v) BSA overnight at 4°C under agitation. Secondary antibodies were diluted to 1/500 in 0.5% (w/v) BSA and incubated 1h at RT under agitation. DAPI (D9542, Sigma-Aldrich) was incubated 10 min.

### Zebrafish experiments

#### Zebrafish husbandry

All animals were kept in a dedicated facility (PRECI - UMS 3444, SFR Biosciences) with a controlled environment: 28°C; 14 h/10 h light/dark cycle, as stated by standard procedures [74]. Zebrafish eggs were collected after spawn and incubated at 28°C in E3 medium (5 mM NaCl, 0.17 mM KCl, 0.33 mM CaCl2.2H2O, 0.33 mM MgCl2.6H2O in distilled water, 0.6 µM methylene blue).

All procedures were conducted in accordance with the guidelines of the European Union and French laws and approved by the local animal ethic committee under regulatory of governmental authority (CECCAPP, Comité d’Evaluation Commun au Centre Léon Bérard, à l’Animalerie de transit de l’ENS, au PBES et au laboratoire P4 (n° C2EA15), APAFIS #33958- 2021100412005226 v6).

#### Toxicity test on zebrafish embryos

Wild-type AB/Tü and Tg(*fli1a*:eGFP)/Casper zebrafish were used. When required, dechorionation was performed with forceps under a stereomicroscope and dechorionated embryos were placed onto a 1% (w/v) agarose gel into the bottom of the wells to prevent them from sticking to the plastic. SEP at different concentrations was added in the medium of embryos at approximately 6-8 hpf. All embryos were monitored at least once a day and dead zebrafish were removed as soon as detected. Tg(*fli1a*:eGFP)/Casper zebrafish were observed at different times under an epifluorescence microscope to report any developmental issue of the cardiovascular system.

#### Zebrafish injection

Zebrafish embryos were anesthetized by immersion in buffered Tricaine methanesulfonate (MS-222 – pH 7) at 0.2 g/L in E3 medium and injected with molecules diluted in PBS using a microinjector (FemtoJet, Eppendorf, Montesson, France) under a stereomicroscope. Injection conditions are detailed in Table 3. After the injection, embryos were kept at 28°C in E3 medium. For the intravascular injection, SEP was coupled to a fluorophore (tetramethylrhodamine-5- maleimide (94506, Sigma-Aldrich)) so that it could be followed easily *in vivo*. SEP was incubated with the fluorophore in excess overnight on rotation, and then purified by 3 to 4 cycles of ITC as described above. Dextran 70 kDa was used as a negative control and was also coupled to a rhodamine (Rhodamine B-isothiocyanate-labeled Dextran; R9379, Sigma-Aldrich).

**Table 3.** Injection conditions for *in vivo* experiments.

|  | **Inflammation** | **Vascular retention** |
| --- | --- | --- |
| **Line** | Tg(*mpx*:eGFP)/Casper | Tg(*fli1a*:eGFP)/Casper |
| **Age** | 3 dpf | 2 dpf |
| **Volume injected** | 2 nL | 1 nL |
| **Administration** | In the yolk sac | Intravascular, in the duct of Cuvier |
| **Injected molecules** | E. coli (500 CFU)<br>SEP (1 or 3.5 mg/mL) | SEP-rhodamine (500 µg/mL)<br>Dextran-rhodamine 70 kDa (500 µg/mL) |

dpf = days post fertilization

#### Zebrafish imaging

Zebrafish larvae were anesthetized by immersion using buffered Tricaine methanesulfonate (MS-222 – pH 7) at 0.2 g/L in E3 medium before imaging.

### Statistical analysis

Data were analyzed for statistical significance using GraphPad Prism Software (version 8.0.1). Precise test choice is mentioned in the figure legend of corresponding data.

## Acknowledgements

We thank the Zebrafish facility (PRECI) and the Protein Science Facility (PSF) from the Federative Structure of Research BioSciences UAR3444/US8. We greatly thank members of the histology and imaging facility (PrImaTiss) at the LBTI institute. We also thank the CIQLE platform (Federative Structure of Research Santé Lyon Est). The Infrastructure de Recherche décentralisée RMN Très Hauts Champs (IR-RMN-THC) Fr3050 CNRS is gratefully acknowledged for conducting the NMR research.

## Fundings

This work was supported by the Agence National de la Recherche, grant numbers ANR-11- TECS-0016 (“DHERMIC”) and ANR-18-CE18-0001 (“Arterylastic”, www.arterylastic.com).

## Declarations of interest

None

## Author contributions

**M. Hoareau**: investigation, formal analysis, methodology, writing – original draft

**C. Lorion**: investigation, formal analysis

**L. Lecoq**: investigation, formal analysis, methodology, resources, writing – original draft

**A. Berthier**: investigation, methodology

**B. Pierrat**: investigation, formal analysis, methodology, software, writing – original draft

**S. Avril**: validation, supervision

**F. Pirot**: validation, supervision

**P. Sommer**: supervision, funding acquisition

**J. Sohier**: investigation, formal analysis, conceptualization

**E. Lambert**: investigation, formal analysis, conceptualization, methodology, supervision, writing – original draft

**R. Debret**: investigation, formal analysis, conceptualization, methodology, project administration, funding acquisition, writing – original draft

## Abbreviations

ADCL: autosomal dominant cutis laxa
CFU: colony forming unit
EDP: elastin-derived peptide
ELR: elastin-like recombinamer
ETD: electron transfer dissociation
GFP: green fluorescent protein
HLE: human leucocyte elastase
hpf: hours post-fertilization
hpi: hours post-injection
hTE: human tropoelastin
HUVEC: human umbilical vascular endothelial
cell IL: interleukin
ITC: inverse transition cycling
LC-MS/MS: liquid chromatography tandem mass spectrometry
LOX: lysyl oxidase
LPS: lipopolysaccharide
MMP: matrix metalloproteinase
NMR: nuclear magnetic resonance
PBS: phosphate-buffered saline
SEP: synthetic elastic protein
SMC: smooth muscle cell
SVAS: supravalvular aortic stenosis
TBS: tris-buffered saline
TNF: tumor necrosis factor
WBS: Williams-Beuren syndrome
β-APN: β-aminopropionitrile

## Notes

### Competing Interest Statement

The authors have declared no competing interest.

