## Supplementary Figures for "A synthetic elastic protein as molecular prosthetic candidate to strengthen vascular wall elasticity"

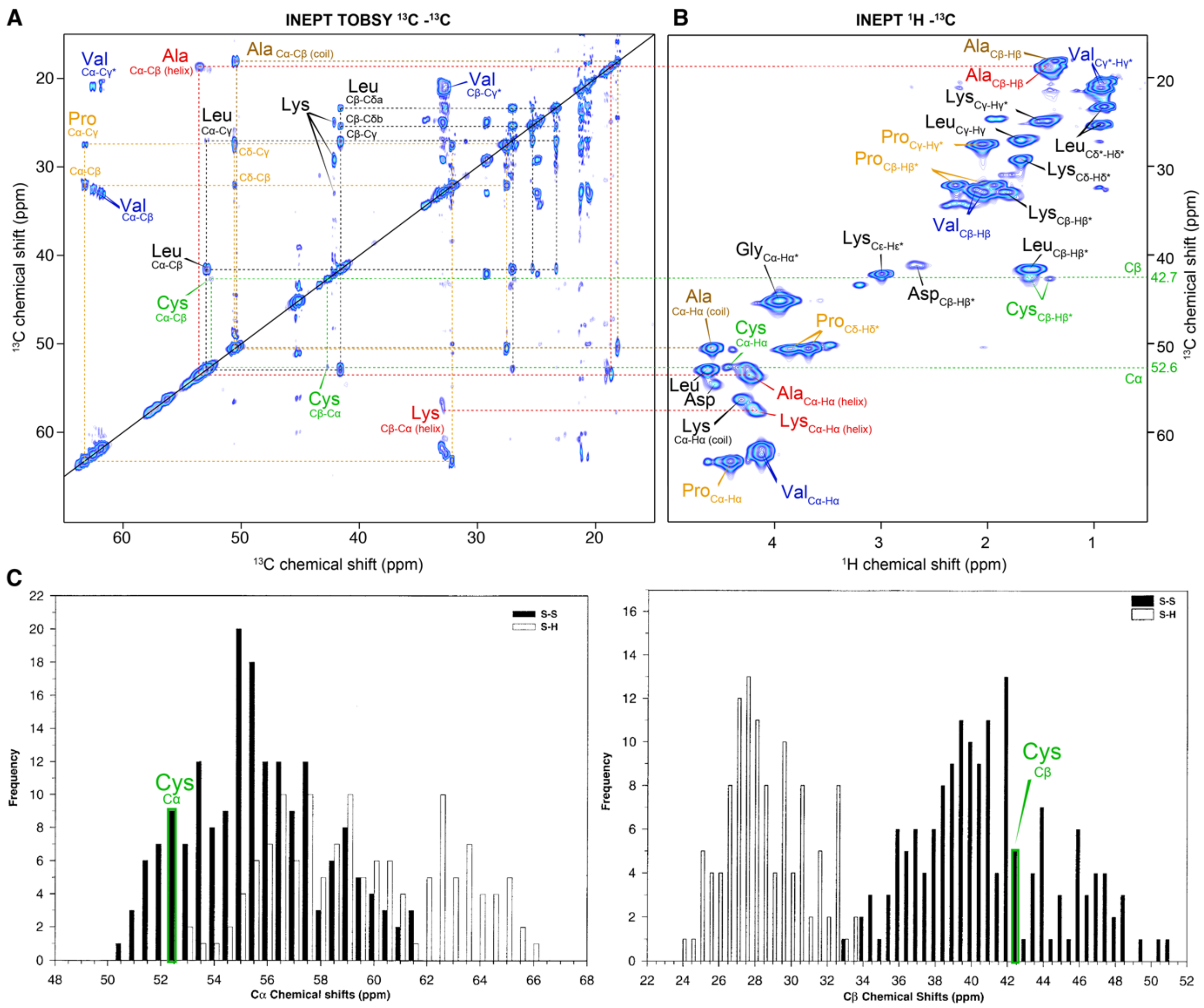


**Fig. S1. C-terminal cystein residues are in oxidized form**

**A)** 2D $^{13}$C-$^{13}$C-INEPT TOBSY spectrum of SEP coacervates, used to correlate carbon chemical shifts of each residue. Dotted lines represent the connected carbon atoms. **B)** 2D $^{1}$H-$^{13}$C-INEPT spectrum of SEP coacervates (also shown in Figure 2C). Dotted lines show the connection with spectrum in A) for the Ala and Lys residues in helical conformation (in red), and for the Cys residues (in green). **C)** Highlight of the chemical shift of the SEP cysteine signal (in green) on the graph of the Cα (left) and Cβ (right) chemical shifts distribution of cysteines, as displayed in [33]. The Cα and Cβ chemical shifts of the cysteine signal in SEP correspond to the oxidized form, suggesting that they form a disulfide bond.

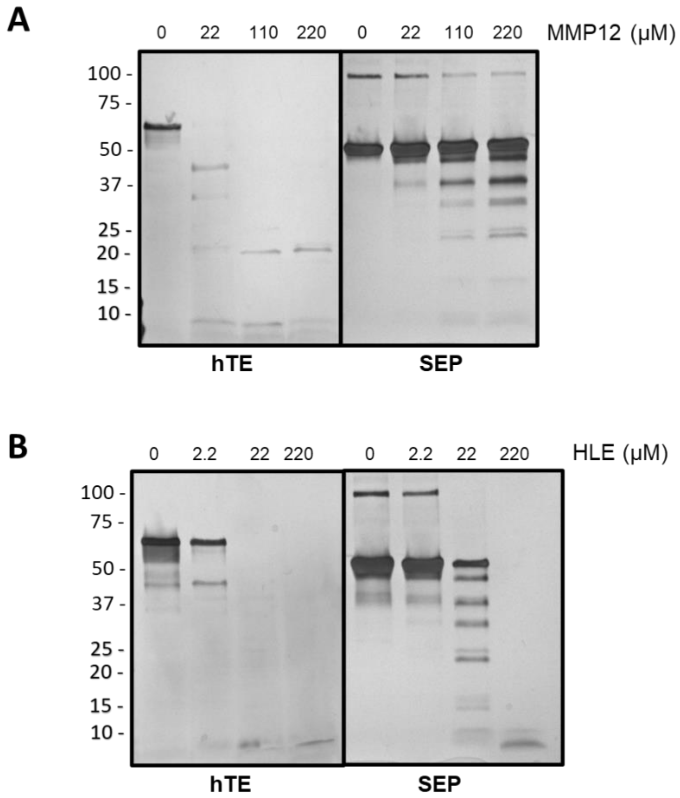


**Fig. S2. SEP is less sensitive to proteolysis by MMP12 and HLE than hTE.**
1 µg of purified SEP or hTE were subjected to enzymatic proteolysis with increasing concentrations of MMP-12 **A)** or HLE **B)** as indicated. Samples were analyzed by silver-stained SDS-PAGE.

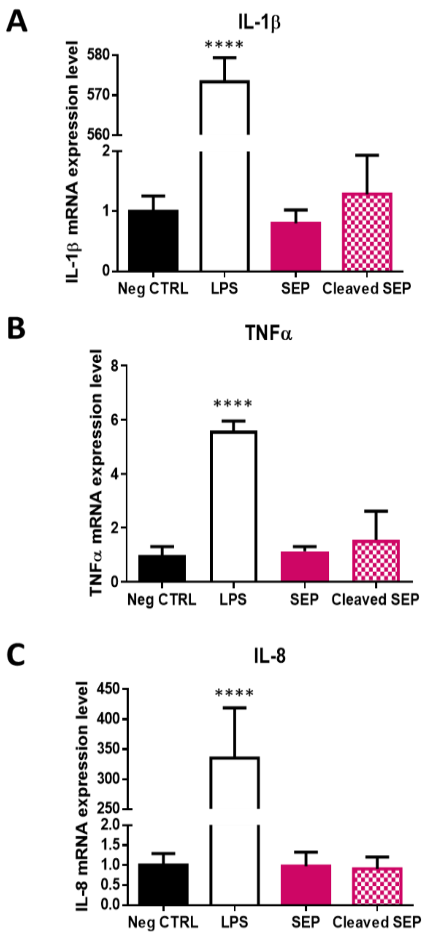


**Fig. S3. SEP or its fragments do not induce pro-inflammatory cytokines expression in THP-1 cells.**

THP-1 cells were treated or not (Neg CTRL) with 1 µg of purified SEP (SEP) or 1 µg of HLE-digested SEP (cleaved SEP) for 24 h prior total RNA extraction. 1 µg of LPS was used as positive control. Il-1β **A)**, TNFα **B)** and IL-8 **C)** mRNA expression levels were assessed by real-time PCR. Statistical analysis by one-way ANOVA and Dunnett's multiple comparisons *post hoc* test, **** $p < 0.0001$.

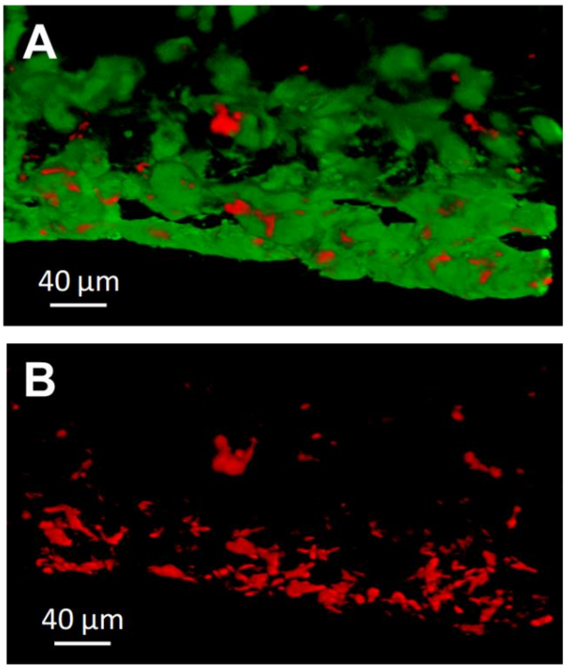


**Fig. S4. SEP integrates blood vessel walls after injection in zebrafish embryos.** Zebrafish embryos Tg(*fli1a*:eGFP)/Casper were injected into the duct of Cuvier with 1 nL of 500 µg/mL SEP at 2 days post fertilization. Confocal imaging was performed 24 hpi on the caudal vein plexus. **A)** Vessels appear in green and SEP was coupled with rhodamine (red) beforehand. **B)** Red channel showing SEP alone. Scale bar = 40 µm.
