## Supplementary Tables for "A synthetic elastic protein as molecular prosthetic candidate to strengthen vascular wall elasticity"

**Table S1:** NMR chemical shifts of residue types present in SEP sequence depending on their secondary structure (helix, coil or beta-strand). Values were taken from the Biological Magnetic Resonance Data Bank (BMRB, https://bmrb.io/).

**BMRB Chemical shifts (ppm)**

| Residue | | Ca | Cb | Ha |
|---|---|---|---|---|
| **Ala** | **Helix** | 55.2 ± 2.8 | 19.7 ± 6.3 | 4.0 ± 0.3 |
| | **Coil** | 52.1 ± 1.9 | 19.3 ± 2.0 | 4.4 ± 0.4 |
| | **Strand** | 51.1 ± 2.1 | 22.9 ± 5.3 | 4.9 ± 0.5 |
| **Arg** | **Helix** | 59.3 ± 2/0 | 29.2 ± 4.0 | 4.0 ± 0.3 |
| | **Coil** | 55.9 ± 2.0 | 31.0 ± 1.9 | 4.5 ± 0.4 |
| | **Strand** | 54.8 ± 2.0 | 33.1 ± 4.5 | 4.8 ± 0.4 |
| **Asp** | **Helix** | 57.1 ± 2.2 | 39.4 ± 5.5 | 4.3 ± 0.3 |
| | **Coil** | 53.8 ± 2.0 | 41.2 ± 1.6 | 4.7 ± 0.4 |
| | **Strand** | 52.9 ± 2.1 | 41.8 ± 3.9 | 5.1 ± 0.4 |
| **Cys-ox (S-S)** | **Helix** | 58.9 ± 3.5 | 37 ± 6 | 4.2 ± 0.6 |
| | **Coil** | 55.3 ± 2.2 | 40.5 ± 4 | 4.85 ± 0.3 |
| | **Strand** | 54.7 ± 2.7 | 45.6 ± 7 | 5.2 ± 0.5 |
| **Cys-red (S-H)** | **Helix** | 62.3 ± 2.6 | 26.2 ± 2.0 | 4.3 ± 0.6 |
| | **Coil** | 58.2 ± 2.6 | 29.4 ± 2.1 | 5.0 ± 0.8 |
| | **Strand** | 56.4 ± 1.9 | 30.2 ± 2.7 | 5.1 ± 0.5 |
| **Gly** | **Helix** | 48.9 ± 4.9 | | 3.8 ± 0.3 |
| | **Coil** | 45.2 ± 1.5 | | 4.1 ± 2.3 |
| | **Strand** | 46.0 ± 0.6 | | 4.2 ± 0.6 |
| **Leu** | **Helix** | 57.7 ± 2.1 | 41 ± 5 | 4.0 ± 0.3 |
| | **Coil** | 54.5 ± 2.0 | 42.5 ± 2.4 | 4.5 ± 0.4 |
| | **Strand** | 53.8 ± 1.9 | 43.4 ± 4 | 4.9 ± 0.4 |
| **Lys** | **Helix** | 58.9 ± 2.0 | 32 ± 5 | 4.0 ± 0.3 |
| | **Coil** | 56.2 ± 1.8 | 32.8 ± 1.7 | 4.4 ± 0.4 |
| | **Strand** | 54.8 ± 1.6 | 35.1 ± 3.6 | 4.9 ± 0.5 |
| **Pro** | **Helix** | / | / | / |
| | **Coil** | 62.6 ± 1.3 | 31.9 ± 1.1 | 4.75 ± 0.4 |
| | **Strand** | / | / | / |
| **Val** | **Helix** | 65.0 ± 3.5 | 31.8 ± 4.3 | 3.6 ± 0.4 |
| | **Coil** | 61.4 ± 2.2 | 32.8 ± 2.0 | 4.3 ± 0.5 |
| | **Strand** | 60.0 ± 2.3 | 34.4 ± 4.1 | 4.7 ± 0.4 |

**Table S2:** NMR chemical shifts of assigned residue types of $^{13}C$-$^{15}N$ SEP, and interpretation about their secondary structure as compared with BMRB values shown in Table S1. Only two residue types show two populations, one in helical and one in coil conformation, namely Ala and Lys residues. In addition, chemical shifts allow to assign the oxidation state of the detected cysteine residues, which are typical of the oxidized form.

| SEP chemical shifts (ppm) | | | | Probable secondary structure |
|---|---|---|---|---|
| | Ca | Cb | Ha | |
| **Ala 1** | 53.2 | 19.0 | 4.23 | helical |
| **Ala 2** | 50.5 | 18.2 | 4.60 | coil |
| **Arg** | 56.0 | 31.0 | / | coil |
| **Asp** | 54.6 | 41.3 | 4.57 | coil |
| **Cys** | 52.6 | 42.7 | 4.42 | ox - coil |
| **Gly** | 45.3 | / | 3.95 | coil |
| **Leu** | 52.8 | 41.9 | 4.64 | coil |
| **Lys 1** | 57.7 | 32.9 | 4.15 | helical |
| **Lys 2** | 56.2 | 33 | 4.33 | coil |
| **Pro** | 63.3 | 32.0 | 4.41 | coil |
| **Val 1** | 62.3 | 32.8 | 4.14 | coil |
| **Val 2** | 61.8 | 33.0 | 4.15 | coil |
